# Image-based phenomic prediction can provide valuable decision support in wheat breeding

**DOI:** 10.1101/2022.09.07.506898

**Authors:** Lukas Roth, Dario Fossati, Patrick Krähenbühl, Achim Walter, Andreas Hund

## Abstract

Traditionally, breeders’ selection decisions in early generations are largely based on visual observations in the field. With the advent of affordable genome sequencing and high-throughput phenotyping technologies, enhancing breeders’ ratings with such information became attractive. In this research, it is hypothesized that G×E interactions of secondary traits (i.e., growth dynamics’ traits) are less complex than those of related target traits (e.g., yield). Thus, phenomic selection (PS) may allow selecting for genotypes with beneficial response-pattern in a defined population of environments. A set of 45 winter wheat varieties was grown at five year-sites and analyzed with linear and factor-analytic (FA) mixed models to estimate G×E interactions of secondary and target traits. The dynamic development of drone-derived plant height, leaf area and tiller density estimations was used to estimate the timing of key stages, quantities at defined time points, and temperature dose-response curve parameters. Most of these secondary traits and grain protein content showed little G×E interactions. In contrast, the modeling of G×E for yield required a FA model with two factors. A trained PS model predicted overall yield performance, yield stability and grain protein content with correlations of 0.43, 0.30 and 0.34. While these accuracies are modest and do not outperform well-trained GS models, PS additionally provided insights into the physiological basis of target traits. An ideotype was identified that potentially avoids the negative pleiotropic effects between yield and protein content.

**Key message:** Genotype-by-environment interactions of secondary traits based on high-throughput field phenotyping are less complex than those of target traits, allowing for a phenomic selection in unreplicated early generation trials.

## 1. Introduction

Like natural evolution, plant breeding is driven by selection. Unlike nature, however, breeders are pressed for time: They have to achieve performance improvements within a few breeding generations, i.e., within a few years. Traditionally, many selection decisions are based on phenotypic observations combined with quality analyses. With the advent of ‘genomics’, these ‘breeders’ eye’ decisions became enhanced with genomic selection (GS) approaches (Meuwissen et al., 2001; Voss-Fels et al., 2019). GS promised to improve selection decisions, in particular for traits where the ‘breeders eye’ has a great potential to fail—e.g., in determining yield potential in single row plots (Rutkoski et al., 2016).

In parallel to genomics research, the field of ‘phenomics’ was growing, increasing both throughput and applicability of phenotyping technologies (Walter et al., 2015). These new phenotyping technologies are seen as key to bridge the gap between lab-determined genotypes and field-observation based phenotypes (Crain et al., 2018). Consequently, secondary traits such as growth dynamics’ traits (Bustos-Korts et al., 2019; Millet et al., 2019; Diepenbrock et al., 2021) or disease resistance traits (Jia and Jannink, 2012) were used as covariates in GS, enabling environment-specific predictions. Nevertheless, for small breeding companies, the entry hurdles for such phenotype-enhanced GS approaches are high, as they are faced with the challenge to simultaneously develop a genomics and phenomics workflow.

Alternatively, it was postulated that phenotyping has also the potential to directly enhance breeders’ decisions by means of ideotype concepts (Donald, 1968). In ideotyp based breeding, one aims to construct a phenotypic ‘model’ (an ideal plant, thus ideotype) representing a unique combination of morphological and physiological attributes to optimize crop performance (Martre et al., 2015). Such a ‘phenomic selection’ (PS)—a selection process for a target trait purely based on (highly processed) secondary phenotypic traits^1^—could be particularly useful in early breeding stages where populations are large and plot sizes small (Rebetzke et al., 2019).

To be successful on the market, breeders have to develop varieties that outperform existing ones in potentially large and inhomogeneous regions, the so-called target population of environments (TPE). The genotype-by-environment (G×E) interaction of target traits such as wheat yield is known to be large, and consequently, breeders must combine small-effect alleles to further increase these traits (Hund et al., 2019). A common set-up in breeding is the use of multi-environment trials (MET) where one can select for a wide adaption of target traits (Smith and Cullis, 2018).

In a recent study, we demonstrated that monitoring soybean METs with high-throughput field phenotyping (HTFP) can reveal strong relations between secondary traits and target traits, allowing to formulate an ideotypes concept (Roth et al., 2022a). Now, this knowledge is transferred to winter wheat (*Triticum aestivum* L.), while additionally including overall performance and stability considerations. It is hypothesized that G×E interactions of secondary traits are less complex than those of the target trait. Under this premise, PS trained on MET data will allow selecting for overall performance and stability even in a single environment—i.e., an early-generation breeding nursery—assisted by high-throughput phenotyping.

A prerequisite of such an approach is the successful implementation of HTFP in METs and breeding experiments. The usefulness of HTFP in breeding is controversial—discussions mainly revolve around the contrast between what *can* be done in phenotyping (i.e., increasing the ‘stamp collection’ of traits) and what is *of value* for breeders (i.e., increasing genetic gain) (Rebetzke et al., 2019). A close interaction of breeders and ‘phenotypers’ is required to avoid the first and achieve the second. The ‘Trait spotting project’ as collaboration between a Swiss plant breeder (Delley Samen und Pflanzen / Agroscope) and the ETH Zurich as academic part aimed to foster such exchange. As part of the project, a drone-based phenotyping platform was developed (Roth et al., 2018b), and methods to extract plant height (Roth and Streit, 2018), canopy cover / leaf area index (Roth et al., 2018a), and tiller count (Roth et al., 2020) for winter wheat were established. Furthermore it was demonstrated that from these low-level traits one can derive three intermediate dynamics’ trait categories, timing of key stages (T), quantities at defined time points or periods (Q), and dose-response curves (C) (Roth et al., 2022c, 2021) (Table 1).

**Table 1:**
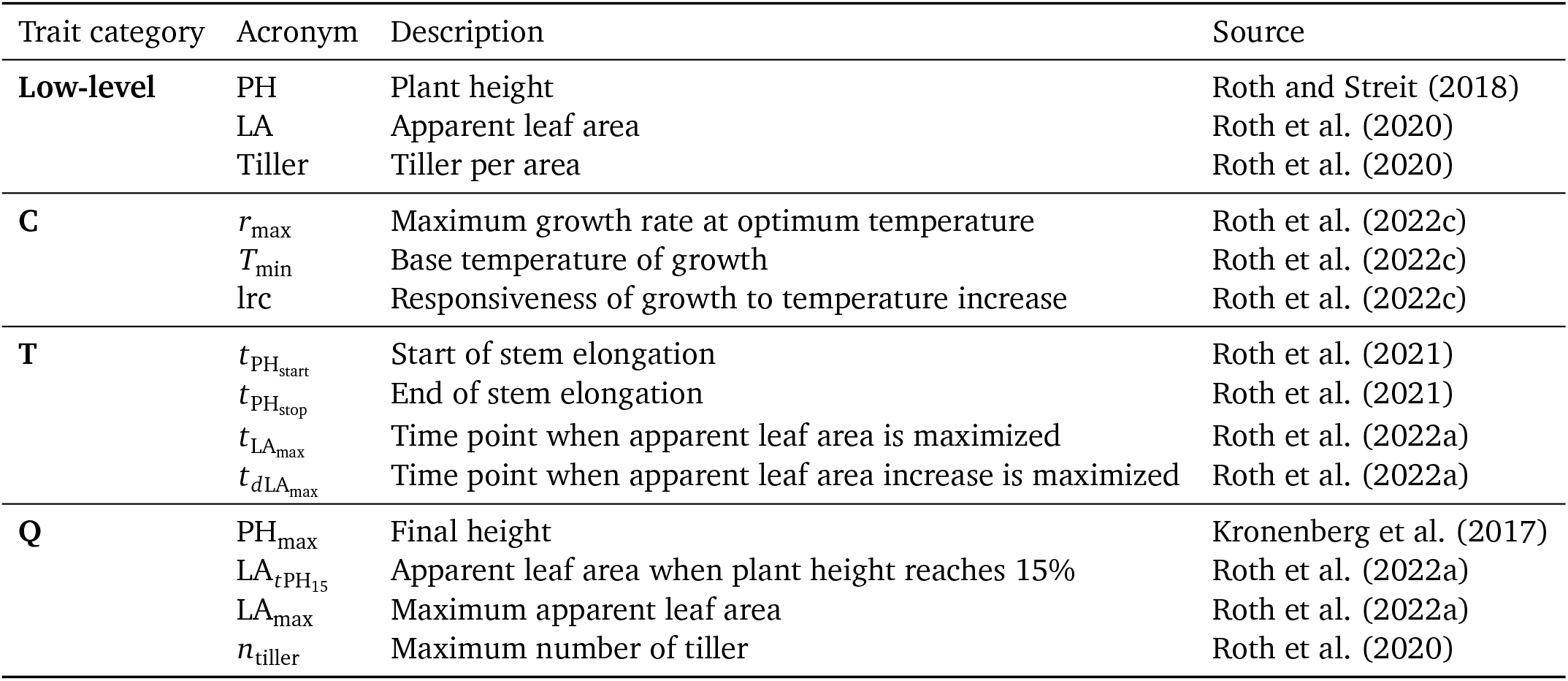
Measured (low-level) traits and processed high-throughput field phenotyping (HTFP) traits of three categories, dose-response curve parameters (C), timing of key stages traits (T), and quantities traits (Q).

With this classification in mind, existing GS and PS literature for wheat focuses mainly on the first two categories. While Rutkoski et al. (2016) and Crain et al. (2018) combined GS with Category Q traits (vegetation indices and canopy temperature measurements at defined growth phases) to predict yield, Sandhu et al. (2021) combined GS with Q traits (vegetation indices at defined time points) to both predict grain yield and protein content. Pure PS approaches are rare to find: Herrera et al. (2018) predicted yield on a spatial grid as result of environmental indices and phenology (Category T). Prey et al. (2020) used spectral indices to predict grain yield and confirmed a significant influence of the measurement time point (Category T).

While these results are of high value for crop physiologists and ecologists to broaden their research, they leave the main question of breeders unanswered: How well can these trait categories be used to complement the selection decision of a breeder, and in what way are they complementary to the ‘breeder’s eye’? This research aims to answer this question on the case example of a prediction model trained on several year-sites of a variety testing experiment and applied to a single-row breeding experiment.

## 2. Materials and Methods

### 2.1. Field experiments

An variety testing experiment (‘Leistungspruefung’/LP01) was performed in two consecutive years (2019, 2020) at three respectively two sites, resulting in a total of five year-site combinations. Respective sites were the field phenotyping platform site of ETH Zurich ‘FIP’ (Kirchgessner et al., 2017) (Lindau Eschikon; Switzer-land; 47.449 N, 8.682 E; 556 m a.s.l., managed according to best practice in Swiss agriculture (‘ÖLN’)); the plant breeding site of Delley Samen und Pflanzen AG ‘Delley’ (Delley, Switzerland; 46.918 N, 6.979 E; 500 m a.s. l., extensively managed according to best practice for growth-regulator and fungicide-free Swiss agriculture (‘Extenso’)); and the testing site of the Strickhof competence center for food and agriculture ‘Strickhof’ (Lindau Eschikon; Switzerland; 47.445 N,8.678, 530 m a.s.l., managed similar to ‘Delley’ (‘Extenso’)).

The experiments were part of the regular testing of advanced breeding material (≥F9) of Agroscope (Nyon, Switzerland)/Delley Samen und Pflanzen AG (Delley, Switzerland) and consisted of 45 elite winter wheat geno-types. Because of the variety testing character, the set for 2019 differed from the set for 2020 by nine genotypes, reducing the number of unique genotypes per year-site to 36.

Year-sites consisted of plots (experimental units in a row-range arrangement with spatial coordinates) enriched with block factors and genotypes. All year-sites contained four replications, except FIP 2020 where only three replications were used. Details about the experimental designs, soil, and management can be found in the Supplementary Materials. Meteorological data was obtained from a weather station next to the experimental field (50 m) for the FIP and Strickhof site and from a public Agrometeo weather station (http://www.agrometeo.ch/, Agroscope, Nyon, Switzerland) in proximity (800 m) for the Delley site. Air temperature was recorded 0.1 m above ground (FIP/Strickhof) and 0.05 m above ground (Delley) every ten minutes and averaged per hour. Growing degree days (McMaster and Wilhelm, 1997) for timing of key stage measurements were calculated assuming a base temperature of 0 °C (Porter and Gawith, 1999).

In 2020, an additional subset of 110 ears-to-row plots of F7 generation lines were monitored at Delley. This single ears descendant experiment was part of the last nursery trial before yield trials. For each line 20 head-to-rows were sown at Delley and, in parallel, 10 head-to-rows at Vouvry (Switzerland), a location with very high disease pressure (powdery mildew, yellow and brow rust, septoria and fusarium head blight). The combination of observations in both locations was used to choose the lines for the yield trials sown the next year. Around 250 lines were selected between approximately 1000 candidate’s lines.

In the subsequent year 2021, the selected lines from the F7 generation were cultivated as F8 generation yield plots. Yield was tested at four locations (Changins, Delley, Villars-le-Terroir, Ellighausen) across the Swiss Central Plateau, which can be considered the Swiss wheat belt. The experimental plots covered 7.1 m^2^ (4.75 × 1.50 m) and consisted of eight rows with an inter-row distance of 0.16 m. Plots were separated by 1.3 m and sown at a rate of 350 seeds m^-2^. Sowing and harvest took place during the months of October and June to July, respectively. The soils are mostly classified as Cambisols (World Resource Base, FAO). The mean value of four cultivars representing the leading cultivars for each Swiss quality class (TOP, 1 and 2), cv. ‘CH Nara’ (class TOP); cv ‘Montalbano’ (class TOP); cv ‘Hanswin’ (class 1); cv ‘Spontan’, (class 2), were used to calculate the relative yield values.

### 2.2. Manual meausurements

For the variety testing experiment, several manual measurements were performed. Beginning of stem elongation (GS30) was determined at three year-sites (Delley 2019 and 2020, FIP 2019) in two replications by destructively measuring the distance between the basal node and the first extending node for three to five representative plants per time point and plot. When this distance reached ten millimeters, the plant was defined to be in the stem elongation stage (Zadoks et al., 1974). Heading (GS59) was defined as the time point, at which 50 % of the spikes fully emerged from the flag leaf sheath (Meier, 2018) and determined manually at four year-sites (Delley 2019 and 2020, FIP 2019 and 2020). Yield was determined by harvesting plots with a combine harvester (FIP and Strickhof: Nursery- master Elite; Delley: Classic; both Wintersteiger, Ried im Innkreis, Austria). Harvested seeds were dried at 30–35 °C if necessary, and the harvest material pre-cleaned in a stand thresher and weighted. Humidity was determined using a HM-400 grain gage (Harvest Data System, Wintersteiger GmbH, Ried, Austria). Yield was then arithmetically normalized to a water content of 15 %. For the subsequent cleaning in an air separator and the estimation of protein content, plot samples of genotypes were merged to mixed probes. Protein content was estimated using a diode array NIR spectrometer (DA-7200, Perten Instruments (today PerkinElmer, Waltham, USA)).

The breeders’ selection notes for F7 lines were based on observations in the preliminary F6 generation in 2019 (one location, Changins) and the F7 lines described in the previous section in 2020 (two locations, Vouvry and Delley). In the F6, heading time, plant height, Zeleny indices, Thousand-Kernel-Weight (TKW), specific weight, grain hardness and (on a 1–9 scale), lodging, septoria tritici, leaf and stripe rust and grain appearance were measured. In the F7, heading date, plant height, lodging resistance, disease resistance for leaf rust (2 notes) and stripe rust (3 notes) and Septoria tritici (1 note) were measured. The uniformity of the 20 head-to-rows in the F7, stand density, and visual estimations of number of ears and length of ears are additional important criterion also taken into account for the selection.

### 2.3. HTFP measurements

The variety testing experiment and the F7 generation experiment were both monitored with the unmanned aerial system (UAS) platform PhenoFly described in detail in Roth et al. (2020). The UAS captured RGB images with high spatial overlap that were processed using Structure-from-Motion software to digital elevation models and camera exposure positions. The flight height was 28 m, flight speed 1.8 m/s, percent end lap 92 %, and percent side lap 75 %. Specific camera settings (e.g., exposure configurations) can be found in Roth et al. (2020). These settings led to a ground sampling distance of 3 mm, restricted motion blur to ≤5 %, and ensured a GCP recover frequency of >70 % for photos that showed one or more GCPs. For FIP 2019, 41 flights were performed between February 19 and July 10; for FIP 2020, 44 flights between February 12 and July 15; for Strickhof 2019, 20 flights between February 22 and July 8; for Delley 19, 21 flights between February 27 and July 12; for Delley 2020, 20 flights between April 6 and July 8.

After Structure-from-Motion processing, digital elevation models were further processed to plant height (PH) traits as described in Roth et al. (2018a). Apparent leaf area (LA) and tiller counts were extracted from processed multiview ground cover images as described in Roth et al. (2020). Based on these three low-level traits (PH, LA, and tiller counts) dynamics’ traits of the first two categories T and Q were extracted according to Roth et al. (2021) using the P-spline/QMER method for LA and PH and using the GS30 based growth model method described in Roth et al. (2020) for tiller counts (Table 1). Dose-response curve parameters were extracted from PH measurements using high-frequency temperature measurements taken by the local weather station combined with lower-frequency drone-based plant height measurements to fit an asymptotic model (Roth et al., 2022c) (Figure 1a). For an overview, all low-level and intermediate traits and corresponding literature references are listed in Table 1.

**Figure 1:**
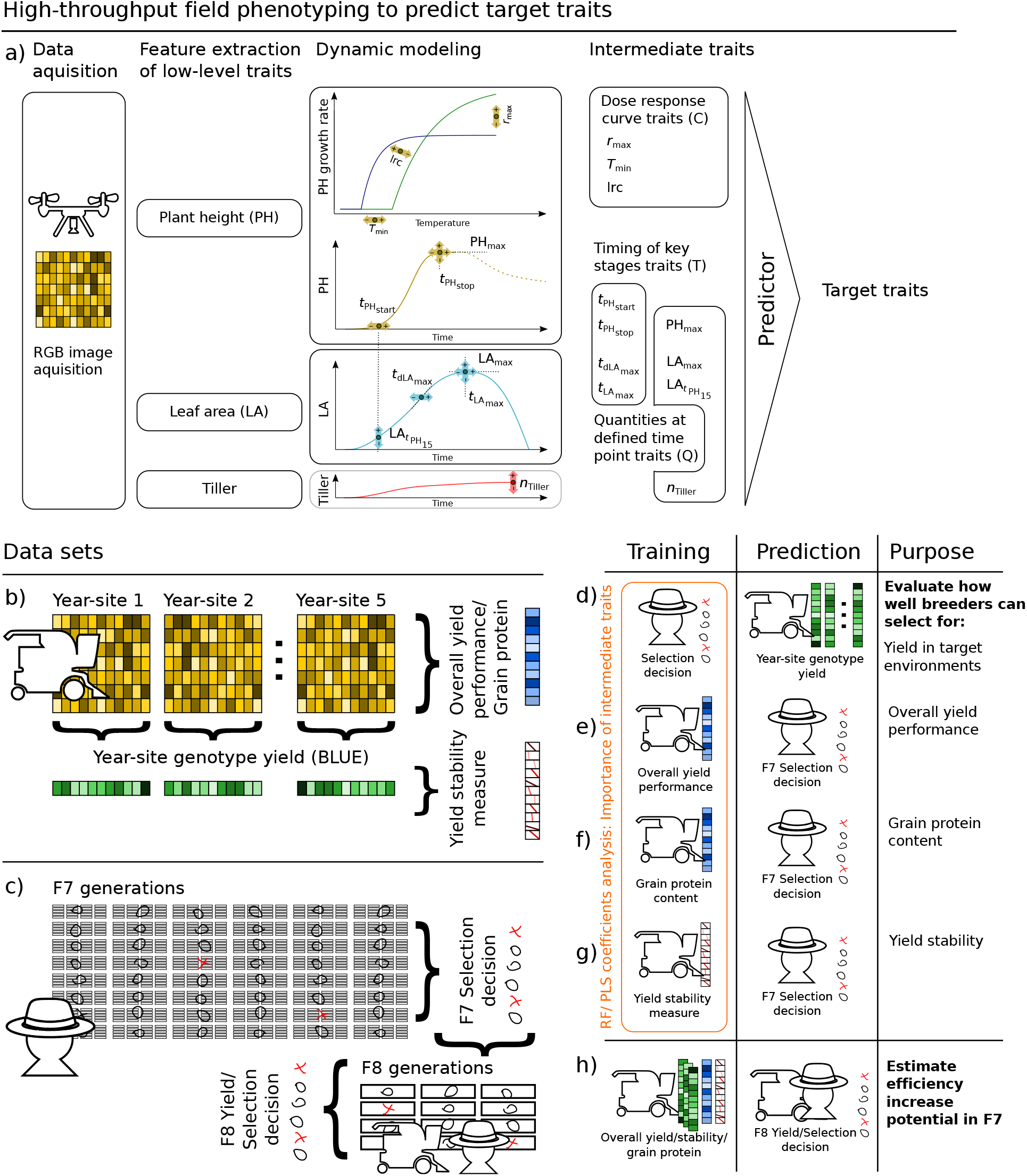
Workflow in high-throughput field phenoptying including data acquisition with drones, extraction of low-level traits, dynamic modeling, and predicting target traits (a), collected data for the variety testing (b) and F7/F8 breeding experiment (c), and how those data sets where used for training and prediction purposes (d–h).

### 2.4. Statistics

#### 2.4.1. Adjusted genotype means per year-site and repeatability calculation

Intermediate traits of all three categories (C, T, Q) were processed in a stage-wise linear mixed model analysis, where the first stage averaged over within-year-site effects and the second stage over between-year-site effects (Roth et al., 2021). For the first stage, the R package SpATS (Rodríguez-Álvarez et al., 2018) was parametrized with the model

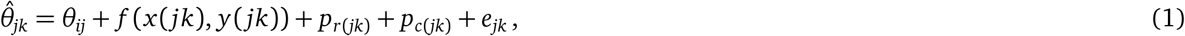

where *i* = (1, …, 45) is the *i*th genotype, *k* = (1, …, 144) the *k*th plot, *j* = (1, …, 5) the *j*th year-site, 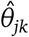 are plot responses based on dynamic modeling, *θ*_*ij*_ year-site genotype responses, *p*_*c*(*jk*)_ range numbers of plots (main working direction, e.g. for sowing), *p*_*r*(*jk*)_ row numbers of plots (orthogonal to main working direction), *f* (*x* (*jk*), *y*(*jk*)) a smooth bivariate surface in spatial *x* and *y* coordinates consisting of a bivariate polynomial and a smooth part (for details see Rodríguez-Álvarez et al., 2018), and *e*_*jk*_ plot residuals with var(*e*) = *σ*^2^*w*^−1^, while *w* are weights based on the standard error estimations from the previous dynamic modeling step, and *σ*^2^ the residual variance parameter. For best linear unbiased estimators (BLUEs) calculations, *θ*_*ij*_ was set as fixed, all other terms as random. For repeatability calculations, *θ*_*ij*_ was set as random. Within-year heritability (repeatability) was calculated according to Oakey et al. (2006).

#### 2.4.2. Overall genotype means and heritability calculation

To calculate overall best linear unbiased predictors (BLUPs) and overall heritability for the five year-sites, a second stage of processing to overall adjusted genotype means (genotypic marginal means) was performed with the R package ASReml-R (Butler, 2018) and the model

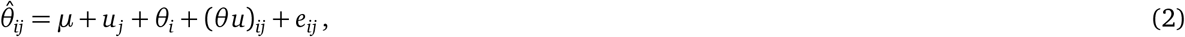

where 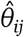 are adjusted year-site genotype means (BLUE) from stage 1, *μ* a global intercept, *u*_*j*_ year-site intercepts, *θ*_*i*_ genotype responses, (*θu*)_*ij*_ genotype year-site interactions, and *e*_*ij*_ residuals with var(*e*) = *σ*^2^*w*^−1^, where *w* are weights based on the diagonal of the variance-covariance matrix from the previous stage, and *σ*^2^ the residual variance parameter. *μ* and *u*_*j*_ were set as fixed and all other terms as random. Heritability was calculated according to Cullis et al. (2006). Genotype year-site interactions (*θu*)_*ij*_ were modeled using a heterogeneous variance model for year-sites (‘Diag’, allowing for both positive and negative variances), and alternatively using an uniform variance model (‘Id’). The model selection was then made based on the Bayesian information criterion (BIC).

#### 2.4.3. Genetic correlation

For the genetic correlation calculation, the univariate model of Equation 2 was extended to a bivariate model (Wright, 1998; Holland et al., 2001),

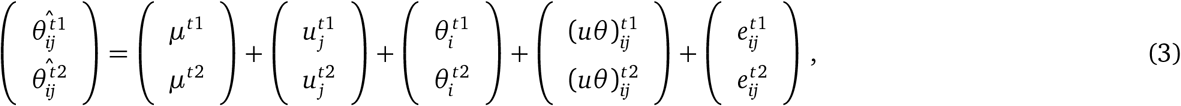

where 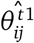 and 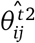 are adjusted year-site genotype means (BLUEs) per trait (trait 1 (^*t*1^) and trait 2 (^*t*2^)), *μ*^*t*1^ and *μ*^*t*2^ global intercepts per trait, 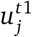 and 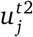 year-site effects per trait, 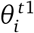 and 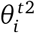 genotype responses, and (*uθ* ^*t*1^)_*ij*_ and (*uθ* ^*t*2^)_*ij*_ the genotype responses to year-site interactions per trait. The terms 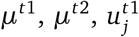 and 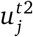 were set to fixed, all other terms to random. Note that *e* and (*uθ*) are confounded, wherefore the two terms were summarized in one variance-covariance structure. Genetic correlations among traits were then calculated based on the estimated variance and covariance components (Holland et al., 2001),

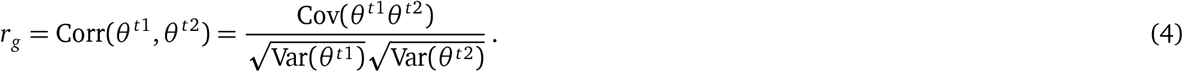

#### 2.4.4. Factor-analytic mixed model for genotype year-site interaction analysis and overall performance and stability calculation

To analyze the severity of genotype year-site interactions for individual traits, Equation 2 was extended with a factor-analytic (FA) component for year-site loadings and genotype scores based on Smith et al. (2001) with a reduced rank (RR) variance model (Thompson et al., 2003),

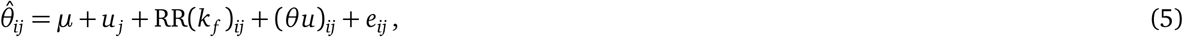

where *k*_*f*_ is the number of factors tested for, *μ* a global intercept, *u*_*j*_ year-site intercepts, and (*θu*)_*ij*_ lack-of-fit effects for genotype year-site interactions modeled using a heterogeneous variance model for year-sites (Diag). This FA model was applied in ASReml-R to all intermediate traits and to the target traits yield and grain protein content. The five year-sites used in this study allowed for a maximum of two factors for the FA model, *k*_*f*_ = (1, 2). Note that instead of the one-stage approach proposed by Smith et al. (2001), a two-stage approach based on year-site BLUEs (Equation 1) was used with residuals *e* set to var(*e*) = *σ*^2^*w*^−1^, where *w* are weights based on the diagonal of the variance-covariance matrix from Stage 1, and *σ*^2^ the residual variance parameter (Piepho et al., 2012).

To determine whether an FA model should be preferred to the simpler linear mixed model (Equation 1) and to determine the appropriate number of factors, two models differing only in the number of factors (FA1–FA2) and two based on Equation 1 were fitted to data, and the performance compared based on the Bayesian information criterion (BIC) and Akaike information criterion (AIC). As the FA models and the linear mixed model differed in the number of fixed factors, the AIC and BIC was based on full log-likelihoods at the REML estimates (Verbyla, 2019) with the function *icREML* provided therein.

The model selection for traits was done based on separate rules for intermediate traits and target traits. For intermediate traits, the ones with complex G×E interactions were discarded from further analyis. For this purpose, traits where the BIC suggested a FA1 or FA2 model were excluded. As this affected only one trait 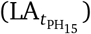, in a second iteration, traits where the AIC suggested an FA1 model with mixed positive and negative loadings were excluded as well 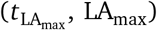, as mixed positive and negative loadings are indicators of crossover G×E interactions. For target traits, discarding traits with complex G×E interactions was not an option, as predicting both target traits was among the aims of this work. Therefore, if both AIC and BIC suggested linear mixed models (Diag or Id), the model suggested by BIC was selected, but if one of BIC or AIC suggestd a FA1 or FA2 model, this model was selected to allow a dissection in stability and overall performance of the target trait. Effectively, this was the case for yield but not for grain protein content.

To estimate the corresponding genotype yield performance and stability across year-sites, the FAST approach as described in Smith and Cullis (2018) was chosen that extracts an overall yield performance (OP) indicator and a stability measure (root-mean-squared deviation, RMSD). Beforehand, the estimates of loadings were rotated as proposed by Smith and Cullis (2018) using the R-code provided in Smith et al. (2021).

#### 2.4.5. Target trait prediction

To predict yield and grain protein content, two approaches were tested: Partial least squares regression (PLS) and random forest regression (RF). PLS uses a linear multivariate model to relate a matrix of observable variables ***X*** to a matrix of responses ***Y*** (Wold et al., 2001). The underlying assumption of correlations among ***X*** allows PLS to analyze noisy and co-linear data, which makes it particularly useful for HTFP. RF on the other hand is an ensemble learning method that combines multiple decision trees to one model, making it similarly suited for highly correlated HTFP data but additionally allowing non-linear mappings of ***X*** to ***Y***. PLS was fitted in R (R Core Team, 2019) with the package *pls* (Liland et al., 2021), RF with the package *ranger* (Wright and Ziegler, 2017). To compare the performance of the two alogrithms (PLS and RF), year-site BLUEs of all three categories (C, T, Q) were used as predictors for year-site BLUEs of yield (Figure 1a, c). Cross-validation (CV) included three different resampling set-ups: (1) Unseen environments (unseen E): 5-fold CV with each fold excluding a whole year-site for the training set that is used as test set, (2) Unseen genotypes (unseen G): 45-fold CV with each fold excluding a genotype for the training set that is used as test set, (3) Unseen genotypes and unseen environments (unseen G and E): 179-fold CV with each fold excluding a complete year-site and a complete genotype while the excluded genotype in the excluded year-site is used for the test set. For RF, hyperparameter tuning was performed for each resampling step using grid-search in a 10-fold CV for unseen E and unseen G and in a leave-one-out CV for unseen G and E. Tuned parameters included the number of trees (ntree, lower: 100, upper: 1000) and the number of variables randomly sampled at each split (mtry, lower: 1, upper: 11). For PLS, the number of components (ncomp) was determined beforehand using all data points in a 10-fold CV per trait.

Both algorithm were then taken to compare the suitability of the three trait categories as predictors for OP (Figure 1a, d), yield stability (RMSD) (Figure 1a, g), and selection decision (Figure 1b, e). Models were trained four times: for PLS with timing traits only (f(T)), with quantities at defined time points only (f(Q)), with doseresponse curve parameters only (f(C)), and for PSL as well as RF with all three trait categories (f(T,Q,C)). The importance of features for PLS was extracted based on the summarized scores for all components per features, and for RF based on 50-fold permutation runs per feature.

For the breeders’ selection decision data set, the response is, in contrast to yield, not continuous but categorical. Hence, the selection scale (line selected, sister line selected, line repeated in next generation, sister line repeated in next generation, discarded) was transformed to an ordinal scale (3, 2, 1, 0, -1) that was then treated as regression problem.

Before model fitting, one outlier was removed for the yield data set (LP01) with *n*_tiller_ *>* 8000. For the breeders’ selection decision data set, 354 out of 2200 rows had to be removed because the dose-response curve fitting failed to converge, resulting in missing values for *r*_max_, *T*_min_ and lrc. As optimization metric for PLS and RF, the root-mean-square error (RMSE) was used, and additionally the normalized RMSE (nRMSE) and Spearman’s rank correlation coefficient (*r*_*s*_) provided.

#### 2.4.6. Efficiency of selection

To compare a pure breeders’ selection decision approach with a HTFP enriched selection decision approach, a HTFP overall yield performance prediction based threshold was evaluated. This threshold was applied to the already performed selection in the F8 generation experiment. The efficiency of selection was then calculated as the ratio between selected lines and total number of lines with and without applying an additional HTFP threshold to breeders’ decisions. Subsequently, the increase in efficiency was derived as the difference between breeders’ selections based efficiency and HTFP enriched efficiency.

## 3. Results

### 3.1. Modeling the dynamic of low-level traits enables to extract heritable but partly correlated intermediate traits

Time series of the low-level trait plant height (PH) indicated a clear start of growth early in the season, followed by a close-to-linear increase phase and a stagnation at a final height afterwards (Figure S1a). While the extracted timing of key stages traits 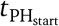 and 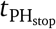 were detected for all plots of the variety testing trials, for the F7 generation breeding trial, the sparse flight density in the early season prevented the extraction of 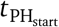 for some plots (Figure S1a, ‘Delley (C), 2020’). The quantity trait PH_max_ was reliably detected for all plots.

Time series of the low-level trait apparent leaf area (LA) showed higher fluctuations between time points and year-sites than PH time series, but also a clear start of growth followed by an exponential growth phase and a decrease phase with high fluctuations afterwards (Figure S1b). Despite that for some plots two peaks of LA were modeled by the P-spline, visual inspections of plots confirmed that the timing trait 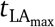 and quantity trait LA_max_ were reliably allocated to the first peak. Consistently, the time point of maximum LA growth 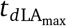 was detected in the increase phase before the first peak (Figure S1b, ‘FIP, 2019’, ‘Strickhof, 2019’). Unlike 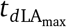, the quantity trait 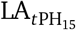 was extracted on a wide range between the start of LA increase and 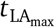.

The extracted dose-response curve parameters for the temperature response of stem elongation showed a large spread of *T*_min_ and *r*_max_ (Figure S1c). While some plots showed almost binary and very unresponsive shapes (Figure S1c, ‘Delley, 2019’), the curves of others appeared more curvy and hence indicated a stronger responsiveness to temperature (Figure S1c, ‘Strickhof, 2019’).

Calculating year-site specific repeatabilities for all intermediate traits revealed strong variations between year-sites for certain traits (Table 2), but no clear trait-independent systematic effect of year-sites. For example, for Delley 2019, the trait lrc showed a below-average repeatability, while other traits were not affected. When comparing HTFP traits with manual measurements, the growth stage ratings GS30 and GS59 showed similar strong variations in repeatability. In contrast, yield repeatability was constant and high for all year-sites. Repeatability values for grain protein content are not available as the measurements were done on mixed probes per genotype. The highest overall heritability was calculated for the growth stage rating GS59, followed by grain protein content and yield (Table 2). For HTFP traits, the heritability varied from very low values (LA_max_) to very high values (PH_max_). Notably, the heritabilities of timing of key stage traits all settled in the middle range (*h*^2^ = 0.52 − 0.71), while the ones for quantities and dose-response curve traits were at the upper and lower extremes.

**Table 2:** Repeatabilities (regular) and heritabilities (**bold**) for intermediate high-throughput field phenotyping (HTFP) traits of the three categories dose-response curve parameters (C), timing of key stages traits (T), and quantities traits (Q) and manually measured growth stage (GS) traits, yield and grain protein content.

| Category | Parameter | Delley, 2019 | Delley, 2020 | FIP, 2019 | FIP, 2020 | Strickhof, 2019 | (all) |
| --- | --- | --- | --- | --- | --- | --- | --- |
| C | $T_{\min}$ | 0.77 | 0.82 | 0.41 | 0.37 | 0.72 | **0.49** |
|  | lrc | 0.18 | 0.82 | 0.22 | 0.25 | 0.57 | **0.84** |
| | $r_{\max}$ | 0.75 | 0.53 | 0.26 | 0.39 | 0.78 | **0.44** |
| T | $t_{\text{PH}_{\text{start}}}$ | 0.44 | 0.32 | 0.70 | 0.01 | 0.91 | **0.71** |
| | $t_{\text{PH}_{\text{stop}}}$ | 0.44 | 0.38 | 0.71 | 0.39 | 0.80 | **0.66** |
| | $t_{\text{dLA}_{\max}}$ | 0.72 | 0.30 | 0.73 | 0.73 | 0.59 | **0.56** |
| | $t_{\text{LA}_{\max}}$ | 0.77 | 0.66 | 0.85 | 0.82 | 0.74 | **0.52** |
| Q | $\text{PH}_{\max}$ | 0.77 | 0.62 | 0.79 | 0.49 | 0.94 | **0.91** |
| | $\text{LA}_{\max}$ | 0.83 | 0.76 | 0.93 | 0.92 | 0.72 | **0.36** |
| | $n_{\text{tiller}}$ | 0.78 | 0.27 | 0.73 | 0.50 | 0.77 | **0.41** |
| | $\text{LA}_{t\text{PH}_{15}}$ | 0.84 | 0.54 | 0.94 | 0.83 | 0.85 | **0.60** |
| Manual | GS30 | 0.89 | 0.44 | 0.71 | - | - | **0.90** |
|  | GS59 | 0.98 | 0.84 | 0.45 | 0.89 | - | **0.97** |
|  | Yield | 0.92 | 0.94 | 0.94 | 0.93 | 0.95 | **0.92** |
|  | Grain protein | - | - | - | - | - | **0.93** |

When calculating genetic correlations, clear relations between HTFP traits and growth stage ratings became visible (Figure 2): For GS59, a very strong positive correlation was found for 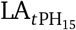 and a strong correlation for *n*_tiller_, indicating that the apparent leaf area in an early growth stage—which was highly (but not significantly) correlated to the number of tillers—is related to the time point of heading. For GS30, a very strong positive correlation to 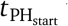 was found, confirming the suitability of 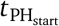 as proxy trait to determine jointing. The partly interdependency of GS30 and GS59 was confirmed by a strong correlation between them.

**Figure 2:**
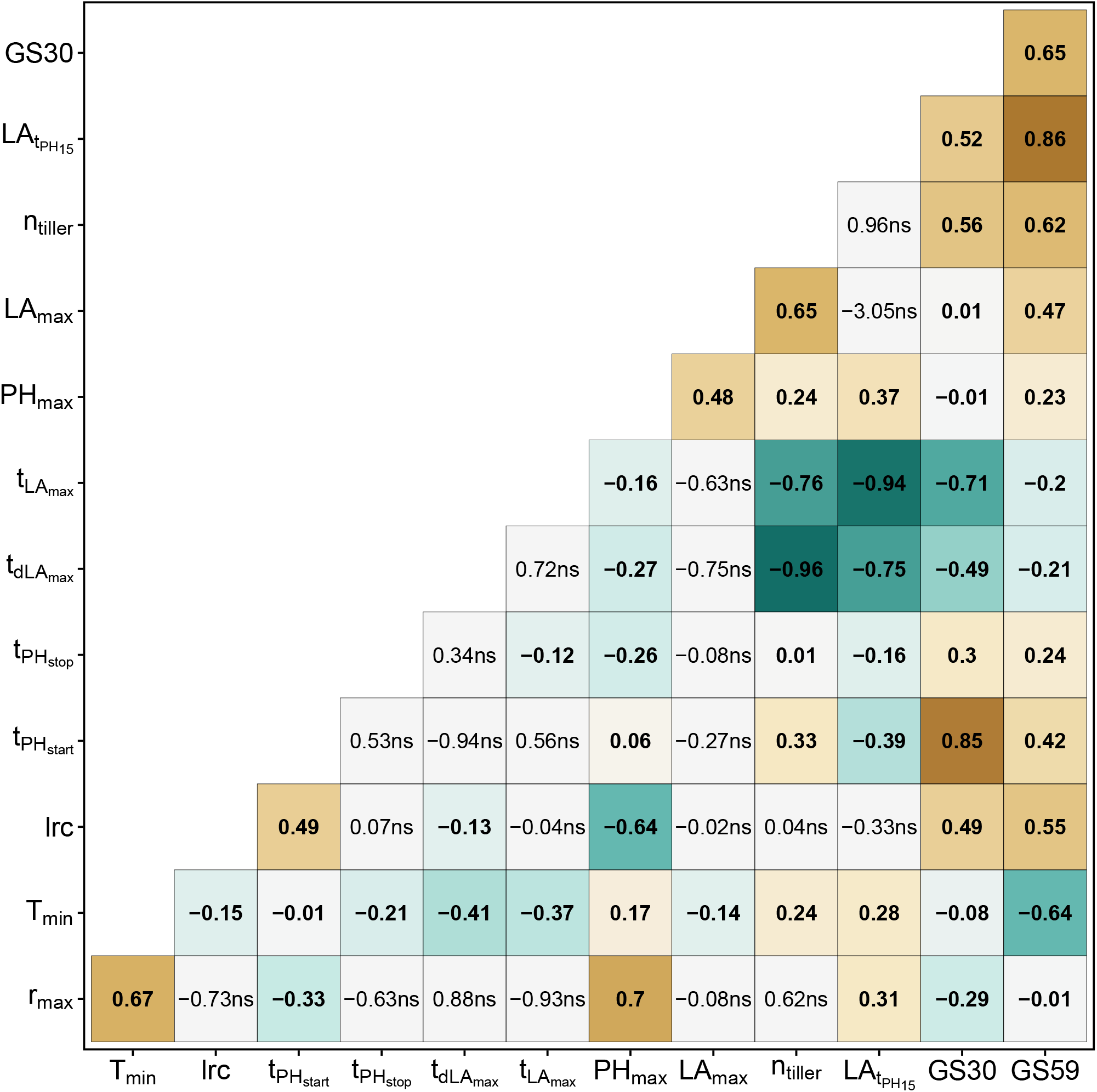
Genetic correlations of year-site BLUEs for HTFP traits and manual growth stage (GS) measurements.

In addition to the previously mentioned relations, a high negative correlation between GS30 and 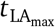 was found, indicating a trade-off between early growth and apparent leaf area mid-season. Nevertheless, a vigorous early growth also led to an early mid-season development, indicated by the strong negative correlation between 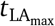 and *n*_tiller_, by the very strong negative correlation between 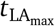 and 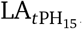, and by the strong negative correlation between 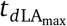 and 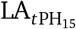. Interestingly, 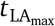 was also negatively correlated with the dose-response curve parameter *T*_min_, indicating that high base temperatures of growth are related to an early time point where the maximum apparent leaf area is reached. This finding was further confirmed by the strong negative correlation between *T*_min_ and GS59.

The strongest relationship was found between 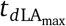 and *n*_tiller_. In addition, *n*_tiller_ was strongly positive correlated to LA_max_. Consequently, high numbers of tillers were associated with an early rapid development of apparent leaf area and large leaf area mid-season.

For the three dose-response curve traits, *T*_min_ and *r*_max_ were strongly correlated. While *r*_max_ and lrc had a strong correlation to PH_max_, the correlation of *T*_min_ with PH_max_ was low, indicating that final height was mainly driven by growth at optimum temperature and steepness of the temperature response. All other relations between HTFP traits showed either only moderate of low correlations or were not significant.

### 3.2. G×E interactions for yield are complex, but less complex for HTFP traits and grain protein content

The courses of meteorological covariates (temperature and precipitation) showed large differences between years and smaller differences between sites (Figure 3d). 2019 was characterized by frequent rainfall throughout the season, but 2020 was characterized by a dry phase between beginning and end of April. Precipitation for the Delley site was higher than for the FIP/Strickhof sites. Temperature courses for all sites were very comparable, but differed between years, with hotter periods towards the end of the season in 2020 than in 2019. Consequently, genotype yields for year-sites (BLUEs) varied between year-sites with lowest yields for Delley, 2020 and highest yields for FIP, 2019 (Figure 3a).

**Figure 3:**
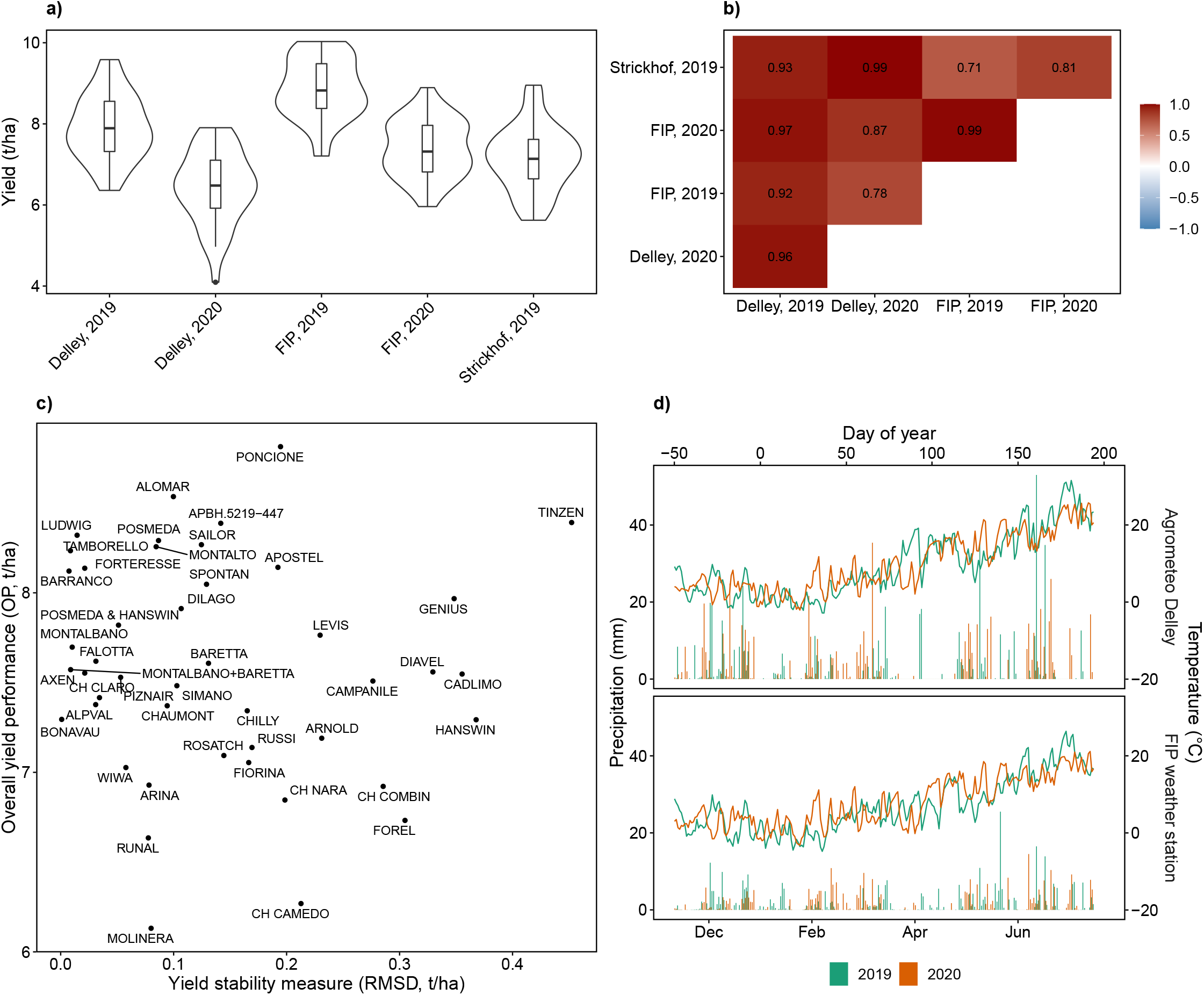
Yield performance of genotypes as BLUEs for individual year-sites (a), estimated genetic correlation for common genotype by environment interactions (b), overall performance versus stability based on a factor-analytic mixed model (FA2) (c), and characterization of sites with temperature and precipitation for 2019 and 2020 (d).

A factor-analytic model with two factors (FA2) showed the best goodness of fit according to AIC but not BIC, partly confirming the assumed high G×E interaction for yield (Table 3). Estimated genetic correlations for common G×E interactions revealed that while some year-sites were highly related (e.g., Strickhof, 2019 and Delley, 2020), others had lower correlations (e.g., FIP, 2019 and Strickhof, 2019) (Figure 3b). Calculating overall yield performance (OP) and yield stability (RMSD) using the FAST approach revealed that the examined genotypes covered well the range of measured OP but clustered at low RMSD (corresponding to high yield stability) with a few outliers with high RMSD (corresponding to low yield stability) (Figure 3c).

**Table 3:** AIC, BIC and full log-likelihood for two linear mixed models with different variance components (Id and Diag) and two factor-analytic models with increasing number of factors (FA1–FA2). Models that did not converge are marked with (-). Lowest AIC and BIC values per intermediate trait are marked in color/*italic*, the selected model in **bold**, * indicate factor-analytic models where the first factor has mixed positive and negative loadings.

| Category<br>Trait | C |  |  | T |  |  |  | Q |  |  |  | Target trait |  |
| --- | --- | --- | --- | --- | --- | --- | --- | --- | --- | --- | --- | --- | --- |
| | $r_{\max}$ | $T_{\min}$ | lrc | $t_{\text{PH}_{\text{start}}}$ | $t_{\text{PH}_{\text{stop}}}$ | $t_{d\text{LA}_{\max}}$ | $t_{\text{LA}_{\max}}$ | $\text{PH}_{\max}$ | $\text{LA}_{\max}$ | $n_{\text{tiller}}$ | $\text{LA}_{t_{\text{PH}_{15}}}$ | Protein | Yield |
| Model BIC |  |  |  |  |  |  |  |  |  |  |  |  |  |
| Id | -635 | 339 | **397** | **1309** | **1450** | 1490 | **1741** | **-1076** | -779 | **2363** | -809 | **85** | 51 |
| Diag | **-662** | **328** | 412 | 1313 | 1461 | **1486** | 1755 | (-) | **-795** | 2366 | (-) | (-) | **45** |
| FA1 | -651 | 342 | 419 | 1322 | 1467 | 1507 | *-1760 | -1066 | *-795 | 2375 | **-845** | 104 | 53 |
| FA2 | -642 | 357 | 432 | 1338 | 1481 | 1521 | 1774 | -1061 | -788 | *2375 | *-839 | 120 | 56 |
| Model AIC |  |  |  |  |  |  |  |  |  |  |  |  |  |
| Id | -657 | 317 | **372** | **1284** | **1425** | 1468 | 1719 | **-1102** | -805 | **2340** | -832 | **63** | 25 |
| Diag | **-700** | **294** | 374 | **1281** | 1425 | **1457** | 1717 | (-) | -814 | 2328 | (-) | (-) | 7 |
| FA1 | -699 | 301 | 374 | 1283 | **1419** | 1466 | * **1712** | -1117 | * **-846** | 2324 | -893 | 69 | 8 |
| FA2 | -699 | 306 | 378 | 1287 | 1423 | 1470 | 1716 | **-1121** | -845 | * **2321** | * **-896** | 72 | **2** |
| Model Log-likelihood |  |  |  |  |  |  |  |  |  |  |  |  |  |
| Id | 335.5 | -151.7 | -177.9 | -634 | -704.3 | -727.1 | -852.5 | 559 | 410.3 | -1163.1 | 422.8 | -24.4 | -4.7 |
| Diag | 362.2 | -135.8 | -175.2 | -630.7 | -701.7 | -719.4 | -846.5 | (-) | 413 | -1151.9 | (-) | (-) | 8.5 |
| FA1 | 364.3 | -137.5 | -173.1 | -629.7 | -694.7 | -719.8 | -841 | 574.6 | 438.9 | -1146 | 461.3 | -23.3 | 9.9 |
| FA2 | 367.3 | -137.1 | -172.2 | -627.6 | -693.7 | -719 | -840 | 579.7 | 440.5 | -1143.4 | 466.2 | -20.9 | 16 |

In contrast to yield, for grain protein content, a simpler linear mixed model with uniform variance for G×E interactions was suggested by both AIC and BIC, indicating only small G×E effects in the examined environments (Table 3).

For all HTFP traits except 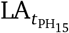, linear mixed models showed better BICs than factor-analytic models, indicating non-complex G×E interactions. Nevertheless, according to AIC, for one timing trait 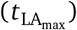 and one quantity trait (LA_max_), factor-analytic models with one factor (FA1) with mixed positive and negative loadings performed best, suggesting crossover G×E interactions. Consequently, 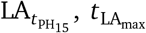, and LA_max_ were excluded from further analysis. Although the AIC also suggested FA1 or FA2 models for 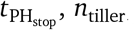, and PH_max_, based on the non-crossover character of the first factor of the FA1 models and the fact that the BIC suggested the most simple linear mixed model with homogeneous variance (Id) for those trait, they were not excluded. Estimations for all remaining traits were based on linear mixed models, depending on the BIC with heterogeneous variance component (Diag) or homogeneous variance component (Id) for genotype year-site interactions.

### 3.3. Partial least square as yield predictor shows less tendency to overfitting than random forest

The performance of predictors for genotype BLUEs with traits of all three categories (C, T, Q) varied strongly depending on the resampling strategy (Table 4). In the most advantageous situation where only the genotype was unseen, RF outperformed PLS with strong correlations and the lowest RMSE and nRMSE. Nevertheless, when using unseen environments as resampling strategy, the correlation dropped for both algorithms, indicating severe overfitting. Further intensifying the resampling strategy to unseen genotypes in unseen environments restored the prediction capacity to some extent for PLS, but less for RF.

**Table 4:** Spearman’s rank correlation (*r*_*s*_), root mean squared error (RMSE) and normalized RMSE (nRMSE) of benchmarks with partial least square (PLS) and random forest (RF) regressors for different resampling strategies and combinations of timing of key stage (T), quantities (Q) and dose-response curve parameter (C) predictors.

| Task | Learner | Resampling | $r_s$ | RMSE | nRMSE |
| --- | --- | --- | --- | --- | --- |
| Yield, Year-site BLUEs |  |  |  | (t/ha) | (%) |
| f(T, Q, C) | PLS (1) | Unseen E | 0.17 | 1.23 | 20.80 |
| f(T, Q, C) | RF | Unseen E | -0.05 | 1.36 | 22.90 |
| f(T, Q, C) | PLS (1) | Unseen G | 0.59 | 0.88 | 14.80 |
| f(T, Q, C) | RF | Unseen G | 0.89 | 0.73 | 12.40 |
| f(T, Q, C) | PLS (1) | Unseen G and E |  | 1.06 | 17.80 |
| f(T, Q, C) | RF | Unseen G and E |  | 1.09 | 18.30 |
| Overall yield performance (OP) |  |  |  | (t/ha) | (%) |
| f(T, Q, C) | PLS (1) | 8-fold rep. cv | 0.43 | 0.49 | 18.20 |
| f(C) | PLS (1) | 8-fold rep. cv | 0.32 | 0.57 | 21.20 |
| f(Q) | PLS (1) | 8-fold rep. cv | 0.40 | 0.50 | 18.70 |
| f(T) | PLS (1) | 8-fold rep. cv | 0.43 | 0.52 | 19.30 |
| Grain protein content |  |  |  | (%) | (%) |
| f(T, Q, C) | PLS (1) | 8-fold rep. cv | 0.27 | 0.74 | 23.90 |
| f(C) | PLS (1) | 8-fold rep. cv | 0.03 | 0.81 | 26.00 |
| f(Q) | PLS (1) | 8-fold rep. cv | 0.34 | 0.72 | 23.20 |
| f(T) | PLS (1) | 8-fold rep. cv | 0.27 | 0.75 | 24.10 |
| Yield stability (RMSD) |  |  |  | (t/ha) | (%) |
| f(T, Q, C) | PLS (1) | 8-fold rep. cv | 0.01 | 0.12 | 26.30 |
| f(T, Q, C) | RF | 8-fold rep. cv | 0.30 | 0.11 | 23.50 |
| Selection decision, F7 |  |  |  | (-) | (%) |
| f(T, Q, C) | PLS (4) | 10-fold rep. cv | 0.26 | 1.21 | 30.30 |

Consequently, PLS was favoured for the target traits overall yield performance, yield stability, grain protein content and breeders’ selection decisions. The hyperparameter of PLS (number of components; ncomp), was retuned for each trait. This resulted in a PLS with one component (PLS(1)) for overall yield performance, stability, and grain protein content, and a PLS with four components (PLS(4)) for selection decision predictions.

### 3.4. Quantity traits are good protein content predictors, overall performance and yield stability predictions profit from traits of all three categories

For overall yield performance (OP) predictions, quantity traits (Q) and timing traits (T) contributed the most to both a high prediction accuracy (*r*_*s*_) and low RMSE (Table 4). Using only dose-response curve traits (C) resulted in poorer predictors. Nevertheless, combining traits of all three categories (C, T, Q) could further improve the RMSE in comparison to using only Q or T traits.

For grain protein content, Q traits contributed the most to a high accuracy and low RMSE. Using only T traits resulted in a slightly lower correlations to grain protein content, using only C traits in a very poor prediction accuracy. In opposite to yield, the performance of a grain protein content predictor that combines traits of all three categories (C, T, Q) was slightly lower than that of a predictor purely based on Q traits. Nevertheless, the combined predictor was favoured for further analysis to allow a comparison of feature scores with other target trait predictors.

For yield stability (RMSD), the prediction performance using PLS was inferior to the accuracy for other traits. Therefore, RF was reconsidered and could indeed restore the prediction accuracy to some extent, indicating that non-linear combinations of traits of all three categories were required to predict stability.

### 3.5. Breeders select for overall performance and yield stability but select against grain protein content

When predicting the breeders’ selections for individual year-site BLUEs, the performance was varying strongly (Figure 4a). Clearly, breeders decisions were more related to a performance in an ‘optimal’ environment than to a low-yielding year-site. Keeping in mind that the F7 generation breeding experiment was actually performed in a low-yielding year-site (Delley, 2020), this capacity for abstraction is impressive: The prediction accuracy was highest for Delley, 2019, a year-site with high yield (Figure 3a) and strong genetic correlations for common G×E interactions with other year-sites (Figure 3b). The lowest prediction accuracy was found for the year-site Delley, 2020, a year-site with below-average yield and lower genetic correlations for common G×E interactions to other year-sites. When predicting breeders’ selections based on overall HTFP BLUPs with measured OP, the predictor performed better than for the best year-site (Figure 4b).

**Figure 4:**
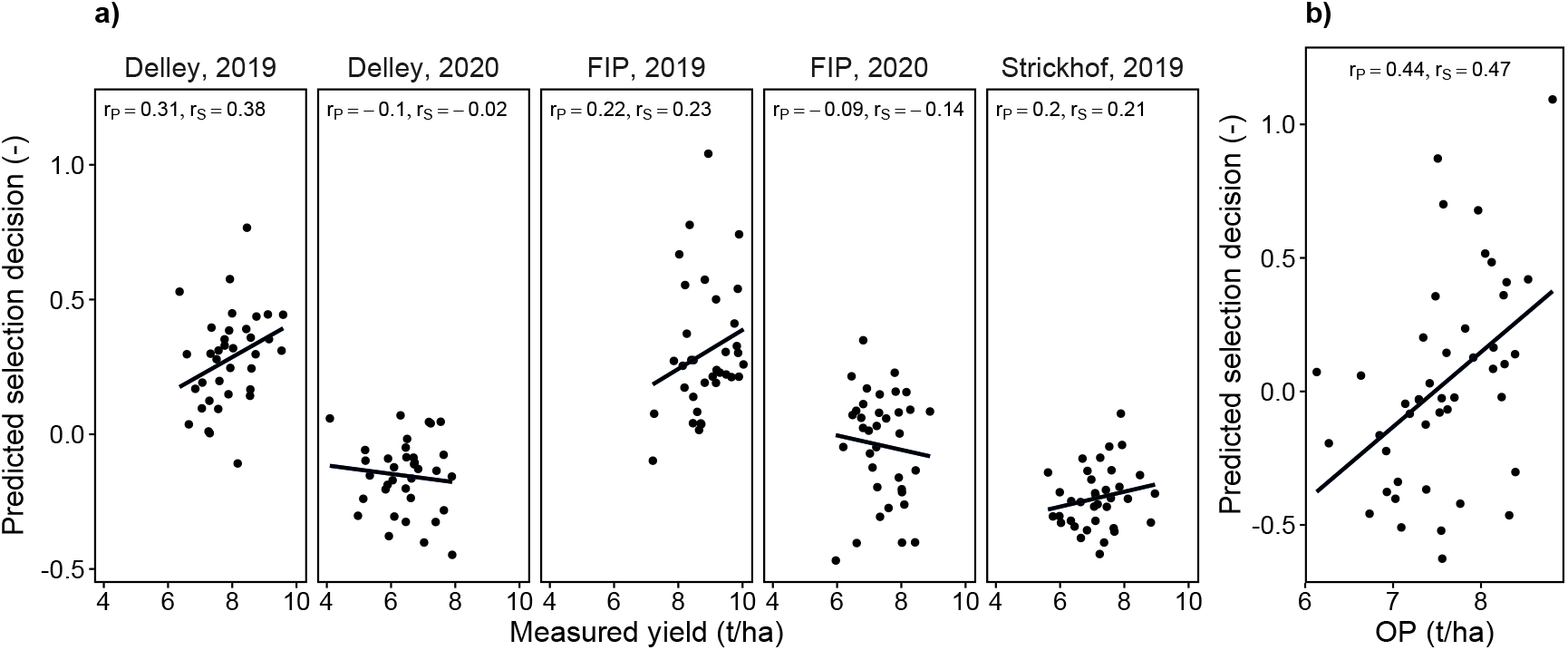
Prediction of selection decision for individual year-sites versus measured yield (a) and prediction of selection decision versus measured overall performance (OP) (b). Pearsons’ correlations (*r*_*p*_) and Spearmans’ rank correlations (*r*_*s*_) are provided, the lines denote a linear regression fit.

The consensus of OP predictions and breeders’ selections became further visible when predicting OP on single lines of the breeding experiment (Figure 5a). The median predicted OP of discarded lines was clearly lower than the one for lines that were selected in the F7 and F8 experiments. Lines that were discarded after F8 were inbetween the two extremes. Breeders avoided lines that had a low OP prediction for their selection. Nevertheless, some lines with above-average OP prediction were discarded.

**Figure 5:**
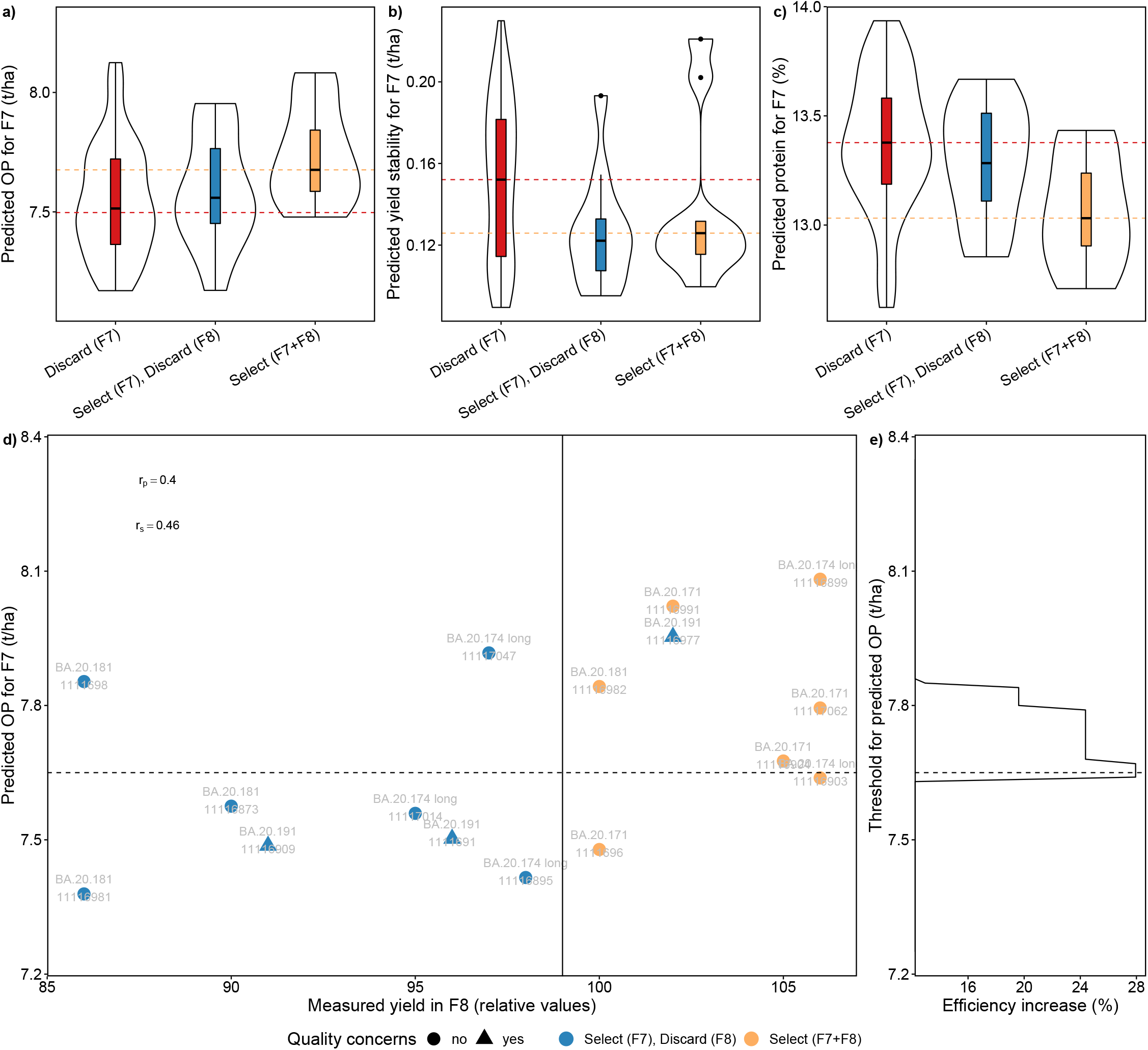
Predictions for F7 single lines for overall performance (OP) (a), yield stability (indicated by a low root-mean-square deviation value) (b) and grain protein content (c), predictions of OP versus measured yield for the subset of F7 lines that were selected and grown on yield plots in the subsequent generation (F8) (d), and efficiency increase if a phenomic selection thresholds would have been applied to the breeding program in F7 (e).

When predicting yield stability (RMSD), medians of selected and discarded lines were close (Figure 5b). Nevertheless, breeders avoided lines with high RMSD in the F7 and F8 experiments, with three exceptions. Again, some lines with above-average stability predictions were discarded.

Predicting grain protein content for single lines in the breeding experiment revealed a lower median grain protein content for selected lines than for discarded lines (Figure 5c). The tendency for low grain protein content was also visible for lines selected in the F7 experiment but discarded in the F8 experiment.

When comparing predicted overall performance with yield measured in the F8 experiment, a clear linear relation for most lines became visible, resulting in an accuracy of *r*_*s*_ = 0.46 (Figure 5d). Applying an overall yield prediction threshold of ∼7.7 t/ha to the F7 selection experiment would have excluded eight genotypes with low yield predictions for the F8. For six genotypes the low yield prediction was confirmed in the F8, but two genotypes would have been excluded by mistake (Figure 5d). In summary, the increase in efficiency if enhancing breeders selection decisions with a HTFP-based prediction threshold would have been in the range of 24–28% (Figure 5e).

Predicting OP, yield stability and grain protein content for the selected F8 lines and F7 lines that were—despite their positive breeders’ ratings—not selected revealed no clear relation of OP versus yield stability (Figure 7a). This result is unexpected, as it is commonly assumed that high-yielding genotypes cannot reach their full potential in every environment, thus decreasing stability. In contrast, the expected antagonistic nature of OP versus grain protein content was confirmed (Figure 7b). Calculating the total harvestable protein per area based on these predictions revealed that selecting for high protein will inevitably reduce the harvestable total protein per area (Figure 7b, grey lines).

**Figure 6:**
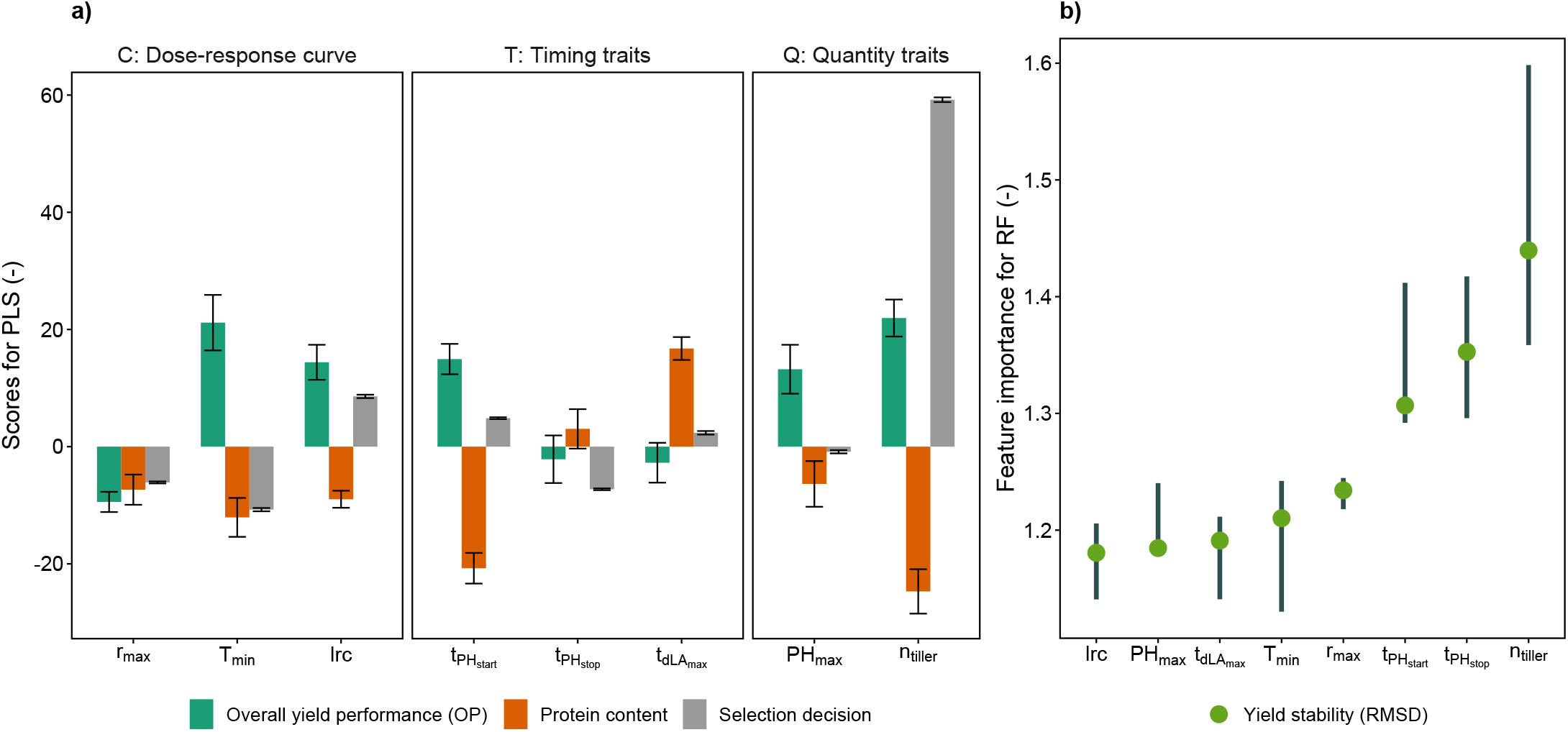
Scores for intermediate HTFP traits for the overall performance (OP) predictor, grain protein content predictor, and breeders’ selections predictors based on partial least squares (PLS) (a) and for yield stability (RMSD) predictor based on random forest (RF). Error bares in (a) are 95% confidence intervals based on jackknife variance estimates, in (b) 95% confidence intervals based on 50-fold cross validation runs.

**Figure 7:**
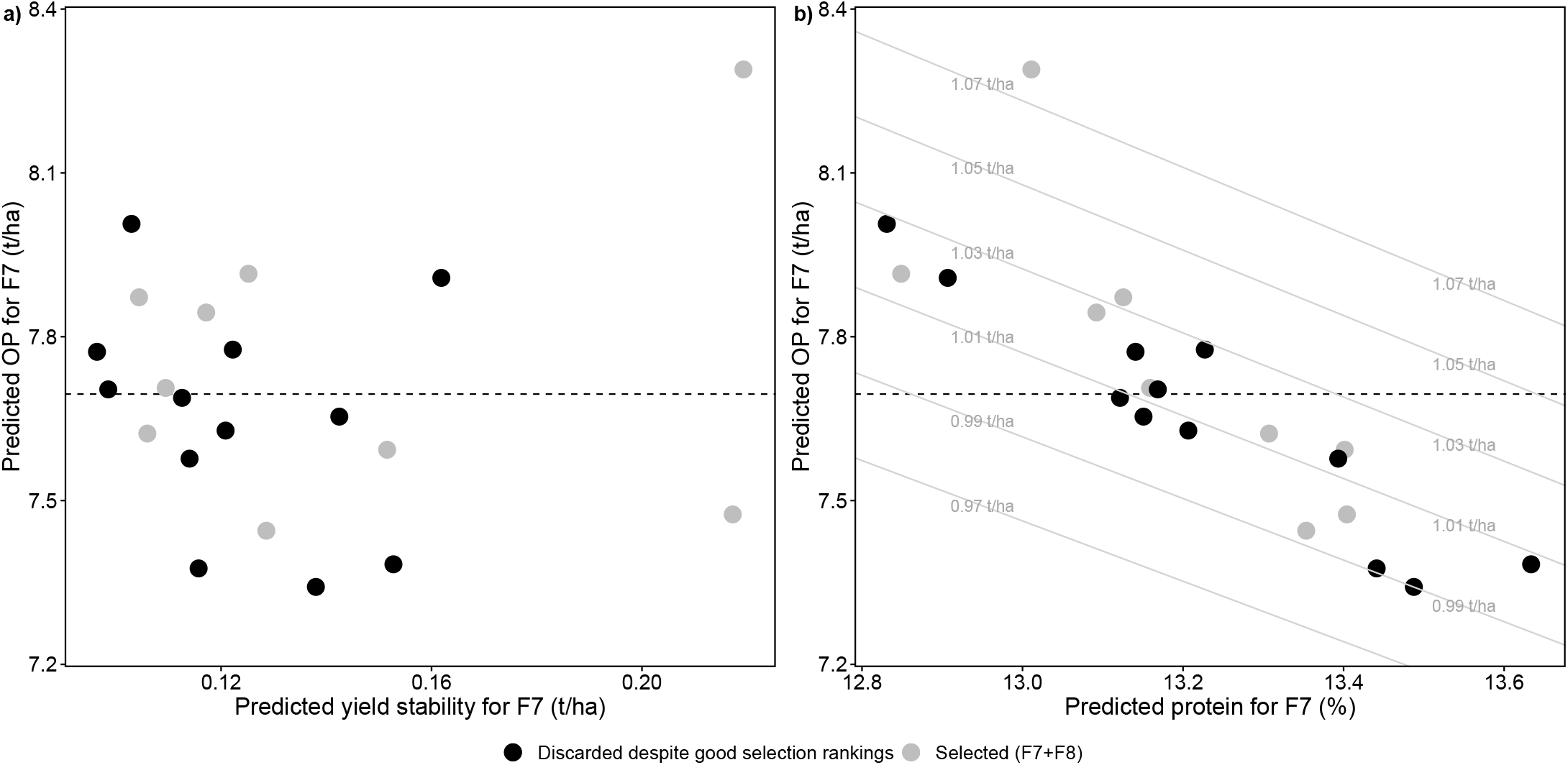
Overall yield performance (OP) predictions versus yield stability (indicated by a low root-mean-square deviation value) predictions (a) and versus grain protein content predictions (b) for the selected F8 lines (grey dots) and for possible candidates proposed by phenomic selection that were not considered (black circles). Numbers on (a) and (b) denote same genotypes, the black dashed line the proposed HTFP selection threshold for OP, grey lines and numbers in (b) denote equidistant protein per hectare lines.

### 3.6. Breeders select differently than HTFP predictors

Extracting PLS scores and RF feature importance of intermediate traits revealed that predictors trained on breeders’ selections and on OP focus on different trait categories and traits. The score for HTFP traits that could explain breeders’ selections was highest and positive for *n*_tiller_, followed by a negative score for *T*_min_, a positive score for lrc, and a negative score for 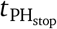. Consequently, traits of all three categories were among the important ones, while the most important one was a quantity trait (Q).

For overall yield performance (OP), the highest score was also found for *n*_tiller_, but the score was less than half the size than for the most important one for the breeders’ selections. The positive score for the quantity trait *n*_tiller_ was shorty followed by a similarly high score for the dose-response curve parameter *T*_min_, for the timing trait 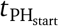, and the dose-response curve parameter lrc. Notably, in opposite to the breeders’ selection, the score for *T*_min_ was positive.

Scores for grain protein content were to a large extent the opposite of scores for OP and/or yield stability. The largest and negative score was found for *n*_tiller_, followed by a negative score for 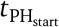.

Feature importance values for yield stability were highest for the quantity trait *n*_tiller_, followed by the two timing traits 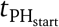 and 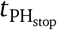. The feature importance of other traits ranged lower, but overall, differences were small, indicating that a non-linear combination of all traits was required to predict yield stability.

Selecting for OP and yield stability simultaneously requires traits that are not antagonistic for those target traits. Such relations were found for the C trait *T*_min_ where the absolute score for protein content was by a factor of 1.8 lower than the absolute score for OP. For the T traits 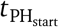 and 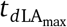 the opposite was observed, having lower absolute scores for OP than for protein content by a factor of 1.4 respectively 6.1. For Q traits, no promising candidates were found.

## 4. Discussion

### 4.1. The suitability of HTFP traits to predict overall yield performance, yield stability and grain protein content

A prerequisite for the indirect selection in breeding experiments is a strong relation of secondary traits to target traits. In this work, it was hypothesized and confirmed that for a subset of the examined HTFP traits, G×E interactions are less complex than for target traits, allowing an efficient selection in early breeding experiments. The low G×E interaction of HTFP intermediate traits may be reasoned by their close direct relation to the mechanistics of growth. In Roth et al. (2021), we elaborated that in particular dose-response curve traits (Category C) are less affected by G×E, as they can be seen as the driver of G×E themselves. Indeed, for the five examined year-sites of this study, all three dose-response curve traits showed simple G×E interactions without crossovers.

In contrast, for two out of four quantity traits (Category Q) more complex G×E interactions were observed, indicating their limited robustness as environment-independent predictors for a target trait. Q traits may be regarded as area-under-the-curve representatives of growth processes (Roth et al., 2021) that are characterized by T and C traits. G×E may increasingly build up during such a growth process. Consequently, a timing trait (Category T) related to one of these Category Q trait, 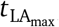, showed more complex G×E effects as well and had to be excluded.

Unlike the intermediate traits, the target trait yield showed partly complex G×E interactions and could be separated in overall yield performance and yield stability. Wheat yield is a trait that can be dissected in physical components, namely, number of plants per square meter, number of shoots with fertile ears per plant, and number and weight of grains per ear (Stern and Kirby, 1979). Each of these components is the result of dynamic growth processes that are driven by genetically determined plant demands and environment supplies (Triboi et al., 2006). Consistently, traits of all three categories were useful predictors of overall yield performance.

For grain protein content, early-season Q and T traits were the most useful predictors. N accumulation in grains is known to be source determined (Martre et al., 2003). Consequently, genotypic variation in grain protein arises mainly from differences in N utilization efficiency and harvest index (Le Gouis et al., 2000). Hence, if N is not a limiting factor, genotypic differences may be very small, and increasing yield will lead to decreased protein content due to dilution. Consistent with this, most intermediate HTFP traits responsible for high protein content were also associated with low overall yield potential. The most influential trait *n*_tiller_ might hint on a dilution by an early development of a high number of ear-bearing culms. To elucidate such effects will require ear counts per unit area, a trait which caught a large interest in the field phenotyping community (David et al., 2021; Dandrifosse et al., 2022).

Additional contrasts for protein content may arise from an adapted growth duration (Triboi et al., 2006). Consistent with this hypothesis, T traits were important as predictors for grain protein content as well. In particular an early start of the generative phase 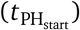 and a delayed maximum growth rate of apparent leaf area timing trait 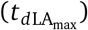 showed high scores for the PLS predictor, reinforcing the finding that changes in phenology were among the main drivers of grain protein content differences.

### 4.2. Formulating an ideotype

As elaborated above, yield and grain protein content are negatively related, to a large part explained by a dilution of protein by accumulated starch of high-yielding varieties. Despite these strong relations between traits, three intermediate traits were identified that allow optimizing for protein content as well as overall performance: From the C traits the base temperature of growth *T*_min_, and from the T traits the time point traits 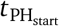 where the stem elongation starts and 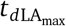 where the maximum increase rate in apparent leaf area is reached.

The most promising ideotype that optimizes overall yield performance and grain protein content requires selecting for high values for *T*_min_ to increase yield, and an early 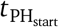 to increase protein (Figure 8a,c). As those two traits are genetically uncorrelated (Figure 2), an independent selection may be easy to achieve. Indeed, in Roth et al. (2022b) we found evidence that for the Swiss varieties such an optimization has already happened. The reported relations of C, T and Q traits of this study are in accordance to the ones reported in Roth et al. (2022b), with one exception: While for the diverse GABI wheat panel examined in Roth et al. (2022b), *T*_min_ was closely related to the phenology traits 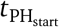 and 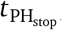, such a relation was not visible for the Swiss elite cultivar set examined in this work. Consequently, the observed effects in the GABI wheat panel may be partly related to population effects, further confirming the possibility for independent selection of *T*_min_ and 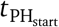.

**Figure 8:**
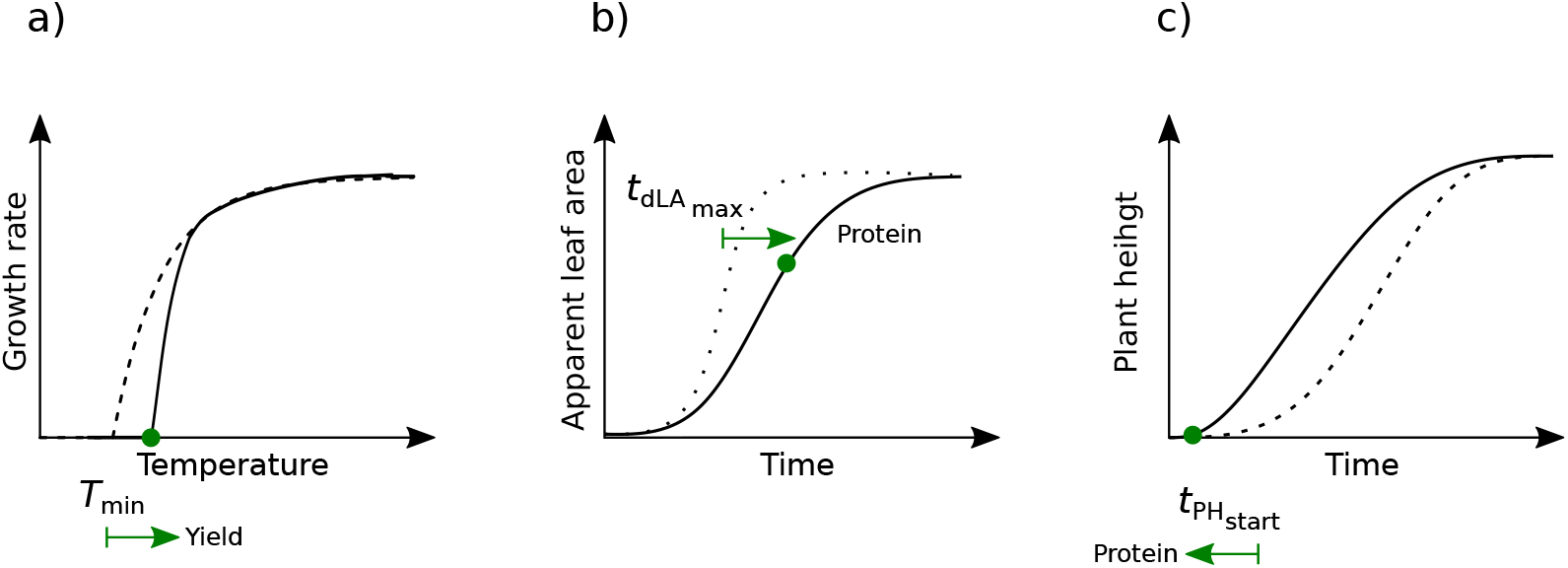
Proposed ideotype to select for high overall yield performance, yield stability and grain protein content based on dose-response curve traits related to the height development of the canopy (a) and timing of key stages traits based on apparent leaf area (b) and plant height (c). The optimization direction of traits are indicated with green arrows.

Alternatively, one could also select for a delayed development of apparent leaf area 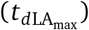 to increase protein content (Figure 8b). Interestingly, a similar ideotype was proposed based on growth dynamics’ HTFP traits in soybean (Roth et al., 2022a). Nevertheless, delaying canopy development in the late season requires special care: In this work, data collection was based on RGB imagery. Delayed T traits for apparent leaf area 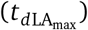 may indicate an advantage of ‘stay-green’ genotypes (Thomas and Smart, 1993). When formulating a corresponding ideotype concept, separating functional from ‘cosmetic’ stay-green is essential—only if canopies remain highly productive in such a stay-green phase a delayed senescence is beneficial. Consequently, if including such traits in a PS approach, one may have to extend the RGB phenotyping technology and include spectral measurements to monitor the dynamics of senescence (Anderegg et al., 2020) and thermal data to extract an indicator for functional stay-green (Anderegg et al., 2021).

### 4.3. Enhancing breeders’ selections with HTFP traits

Training a PLS with breeders’ selection decisions and predicting year-site specific yield revealed that breeders select for an average performance for their TPE. Notably, the training set was based on selection decisions taken on a year-site that showed lower-than-average genetic correlations for common G×E interactions to other yearsites. Consequently, the year-site was unfavourable for breeders as selection base. Nevertheless, the breeders managed to select in a year-site ‘neutral’ manner for overall performance, indicating that their selection was based on robust secondary traits with little G×E interaction.

In opposite to overall yield performance, grain protein content was a trait on which selection does not seem to have a strong effect. While most of the traits showed converse effects on protein content and yield, some had a stronger effect on either of both. While an indirect selection for a high grain-protein deviation based on such information may be difficult, the information how intermediate traits affect these target traits is of major importance for breeders. Early vigor for weed suppression for example is increasingly important due to the political pressure for herbicide-free cropping systems. Thus, an increased vigorous development to achieve a better weed suppression in spring may reduce protein content while increasing yield.

For yield stability, breeders’ selections were neutral, they did not neglect but also not strongly select for more stable genotypes. Again, with HTFP, breeders selection decisions could become enhanced with a stronger focus on stability. However, quantifying yield stability will need a large number of test environments and genotypes. The five year-sites included in the present study offer only a limited insight into the driving factors of G×E. Implementing HTFP in a lager number of sites including future climate sites may greatly enhance our understanding about ideotypes with a strong resilience to extreme weather conditions.

Introducing a threshold based on overall yield performance predictions with HTFP traits could increase the efficiency of selection by up to 28%. Predictions for the discarded lines in the F7 breeding experiment made evident that potential candidates with high yield and protein potential were present in the breeding program.

The prediction accuracy (*r* ≈ 0.4) of PS trained on a small set (45 genotypes) was comparable to those measured in GS approaches with large training sets (*>* 550 genotypes) (Rutkoski et al., 2016; Crain et al., 2018; Sandhu et al., 2021). If keeping all other parameters of a breeding program constant, such an efficiency increase would directly translate into genetic gain, making PS for the particular case as competitive as GS. The achieved rank prediction accuracy of unseen genotypes in an unseen environment of 0.46 further underpins this finding.

In addition, PS and GS do not necessarily have to be seen as competing—integrative approaches may also be conceivable. Diepenbrock et al. (2021) for example used GS to predict parameters of a crop growth model that related to the target trait yield, which improved the accuracy of yield predictions for unseen genotypes in unseen environments. In this work, it was demonstrated that if the TPE is part of the PS training set, the non-complex G×E nature of intermediate HTFP traits may allow comparable target trait prediction approaches without explicitly including environmental covariates. In return, this could potentially allow predicting intermediate traits directly with GS instead of measuring them with HTFP.

Once trained, the presented phenomic selection approach can be applied to single-row plots in early generations. Nevertheless, the training set requires to be collected in generations grown in yield plots. This study was conduced within the framework of the Swiss winter wheat breeding program of Agroscope/Delley Samen und Pflanzen AG, without interfering with it. Starting with the F8 generation, yield plots are used in this breeding program. The F8 generation is grown in one year only, and most of the examined ∼200 genotypes discarded afterwards. An essential fundamental of the presented approach is to examine G×E interactions of target traits (yield, grain protein content) and high-throughput phenotyping traits. As source for G×E interactions in wheat cultivation in Switzerland, the year is as important as the location (Figure 3b). Using the F8 generation as training would have prevented from examining year-based G×E interactions, therefore limiting the power of the training set. Consequently, it was taken advantage of the variety testing character of the subsequent F9–F12 generation experiments that are performed in multiple years. Depending on the breeding program, the generation that is best suited as training and selection set may vary. Importantly, the training set has to be as genetically close the the selection set as possible, while providing sufficient reliable yield measurements in multiple locations and years. For the Swiss winter wheat breeding program, this prerequisite is fulfilled in the F9–F12 generation experiments.

### 4.4. Considerations on the added value of PS versus GS to support the Swiss winter wheat breeding program

The attractiveness of a particular approach for a breeding program depends to a large extent on the costs. In the following, the economical costs and accuracy increase of PS versus GS is evaluated on the case example of the Swiss winter wheat breeding program of Agroscope (Nyon, Switzerland) / Delley Samen und Pflanzen AG (Delley, Switzerland). A base assumption is that the design of the program itself remains unchanged and that GS and PS could be introduced independently at certain breeding stages. Cost estimations for GS are based on know-how gained during the implementation of GS in the Agroscope/Delley Samen und Pflanzen AG breeding program. Cost estimations of PS are base on annual costs of the Trait Spotting project where two sites per year with a total area of 2.160 m^2^ were monitored.

Given is the winter wheat breeding program with 300’000, 12’500, 5’000, 2’500, 900, 220, 60, 30, 36, and 36 genotypes in the F3–F12 generations (Figure 8a). Given is further that F3 are single plant, F4 single rows, F5 double row plots, F6 four-row plots, F7 20-row plots, and F8–F12 yield plots, while rows have a 0.25 × 1 m and yield plots a 1.5 × 5 m area (Figure 8b). Finally, given is that for F3–F6 one location, for F7 two locations, for F8 four locations, and for F9–F12 five locations are used (Figure 8a). The measured accuracy for yield prediction with PS in this work was *r* = 0.43, for measuring yield with yield plots *r* = 0.9, and the provided accuracy for GS in wheat is *r* = 0.32 (Crain et al., 2018).

When using these values to calculate total costs per breeding stage, costs for GS in early breeding stages (F3– F6) where number of required samples are high but the areas to monitor small are higher than for PS (Figure 9b). Starting with F7, GS becomes more affordable than PS. Finally, starting with F8, yield plots could be use, representing the most cost-efficient selection strategy.

**Figure 9:**
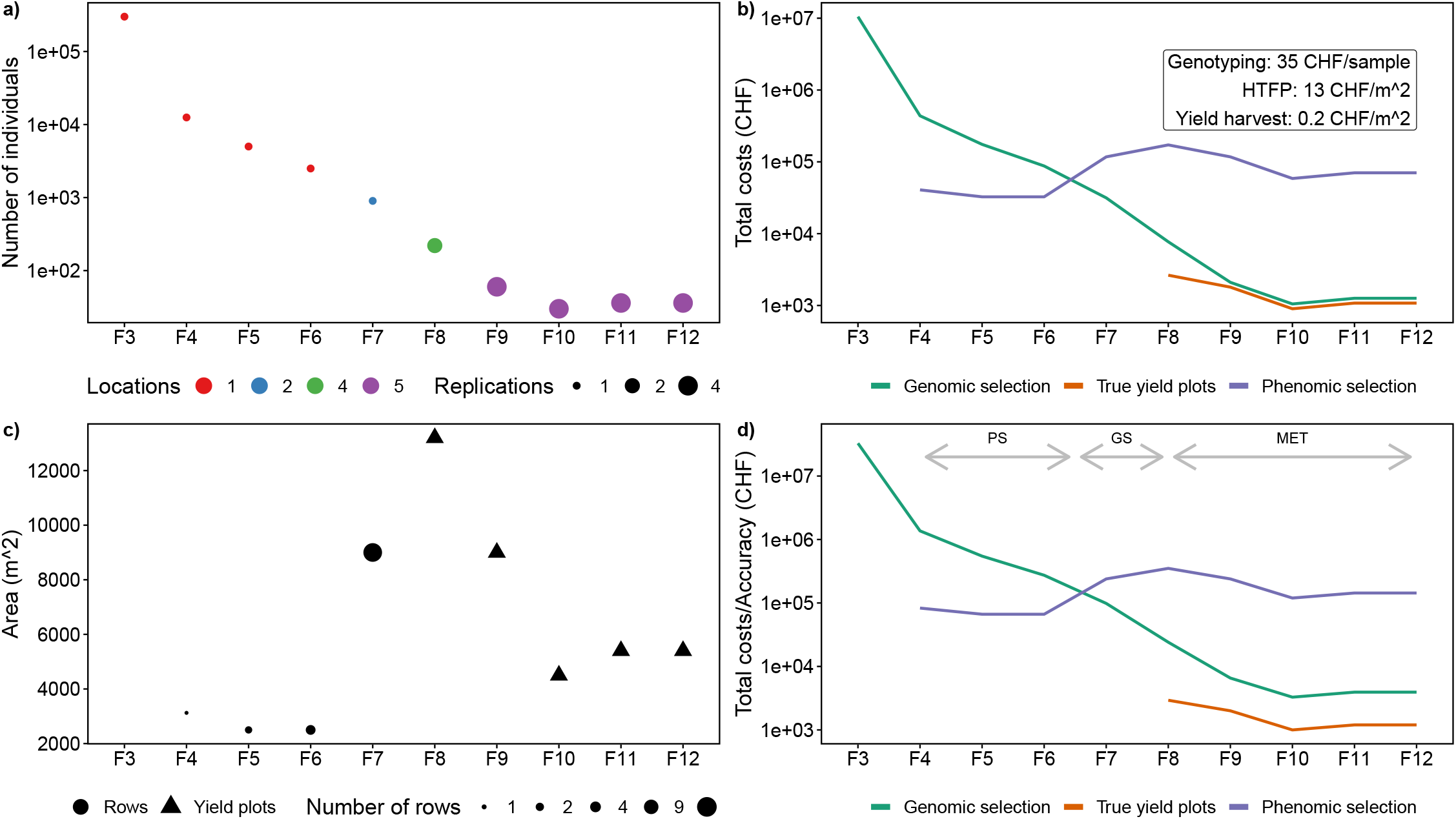
Dimensions of the Swiss winter wheat breeding program (a, c) and costs (b) versus accuracy (d) of a genomic selection approach (GS) based on genotyping and a phenomic selection approach (PS) based on high-throughput field phenotyping (HTFP) and classical yield multi-environment trials (MET).

Given that both GS and PS are not yet fully part of a breeding program yet, breeders would be more interested in the return in accuracy increase if adapting new technologies than in total costs. When dividing the total costs by the expected selection accuracy, costs of increasing the accuracy of selection are lowest for PS in early breeding stages, lowest for GS in intermediate breeding stages, and lowest for yield trials in late breeding stages (Figure 9d). In summary, for the Swiss winter wheat breeding program, a hybrid GS/PS strategy may be best suited to increase the genetic gain, with PS focusing on early single and double-row plots.

## 5. Conclusion

Using HTFP to extract dynamic traits of three different categories (C, T, Q) derived from drone-based RGB imaging proved to be suitable to allow a prediction of overall yield performance, yield stability, and grain protein content. Less complex G×E interactions of HTFP traits than those of target traits allowed the application of HTFP even in early breeding generations with few locations. Traits related to quantities at defined time points or time periods (Category Q) can be seen as proxy traits for yield components, and are the result of related growth processes, giving them high prediction potential for overall yield performance and protein content. Timing of key stages (Category T) and dose-response curve (Category C) traits are more related to stability. This suggests focusing on such traits when aiming at the mitigation of the effect of abiotic stresses. As dose-response curve traits have lowest G×E interactions, they potentially allow performing predictions on HTFP data from single locations.

The prediction accuracy of phenomic selection (PS) is comparable with that reported for genomic selection (GS) and allows increasing the selection efficiency by up to 28%. While costs of GS mainly depend on the number of genotypes to sample and continuously decrease with increasing generations, PS depends on the area to monitor, making it cost-efficient for early breeding stages. Other than GS, PS allows analyzing the influence of dynamic traits on target traits, providing both insights on existing breeders’ selection processes but also formulating an ideotype concept for further refinements of those. In particular the C traits base temperature of growth, and the T traits start of stem elongation and time point of maximum apparent leaf area increase allowed formulating an ideotype concept that may optimize yield, yield stability and protein content.

## Acknowledgement

We would like to thank the staff from Delley Samen und Pflanzen (DSP), namely Karl-Heinz Camp for coordinating the Trait Spotting Project, Flavio Fossati for preparing and exchanging data sets of manual ratings, Etienne Thévoz for field management at the Delley site, and Yves-Etienne Cornamusaz for performing flights at the Delley site. Furthermore, we acknowledge Hansueli Zellweger (ETH Zurich) for field management at the FIP site, Helge Aasen (ETH Zurich) for supervision and support with drones and data processing, and Simon Treier for help with data collection.

## Statements & Declarations

### Author Contributions

L.R: Conceptualization, investigation, methodology, software, formal analysis, visualization, writing—original draft. D.F.: Investigation, Validation, Data Curation, writing— review & editing. P.K.: Investigation, Validation, Data Curation, writing— review & editing. A.W.: Conceptualization, writing— review & editing. A.H.: Conceptualization, supervision, project administration, funding acquisition, writing— review & editing.

### Data Availability

The datasets generated and analyzed during the current study are openly available in the ETH Research Collection repository, http://doi.org/10.3929/ethz-b-000566864. Source code for the phenomics data processing methods used in this study are openly available in the ETH gitlab repository, https://gitlab.ethz.ch/crop_phenotyping/htfp_data_processing.

## Appendix

### Figures

**Figure S1:**
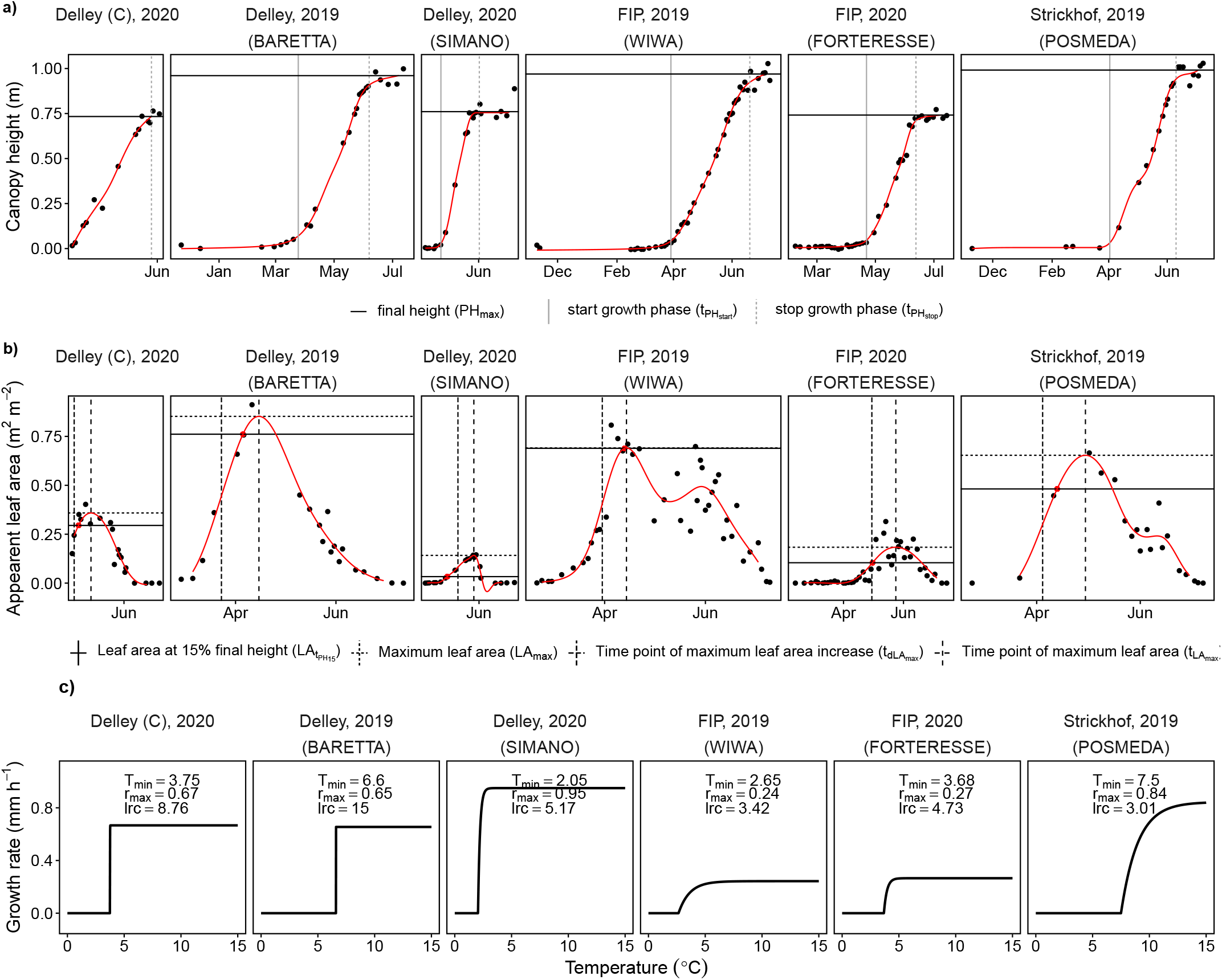
Examples of extracted low-level traits canopy height (a) and apparent leaf area (b) with measured time points (black circles) and fitted P-splines (red lines) and fitted temperature dose-response curves (c) for six plots of six year-sites.

## Site descriptions

### Site Delley

#### Delley19 (Genevey 2)

The site Delley 2019 was located at Delley, Switzerland (46.918 N, 6.979 E, 500 m a.s.l.). The soil had a clay content of 15-25%, humus content was 1.5% and pH 7.2. Soil characteristic were determined in 2017 (Sol Conseil, Nyon, Switzerland). Preliminary to wheat, pea (*Pisum sativum* L.) were grown. After preliminary crops were harvested, the soil was irrigated with 20 mm water, plowed and harrowed before wheat was drill-sown.

For the design, varieties were arranged in a 6 × 6 lattice design with four replications. The experimental unit was a plot of 9 m^2^ (1.5 m in row and 6 m in range direction including paths and wheal tracks), resulting in a harvested surface of 7.05 m^2^.

Wheat was drill-sown in 8 rows per plot with a row distance of 0.125 m on October 24, 2018. Sowing density was 350 plants m^-2^.

On February 25 and April 1, 84 kg N ^-1^ and 65 kg N ^-1^ ha^-1^ were applied. Additionally, in total 50 kg phosphorous ha^-1^, 75 kg potassium ha^-1^, 20 kg calcium ha^-1^ and 18 kg magnesium ha^-1^ were applied.

#### Delley20 (Grandcour)

The site Delley 2020 was located at Gletterens, Switzerland (46.88971 N, 6.93694 E, 482 m a.s.l.).

Soil characteristics: Clay content: 20%, pH: 6.4, humus content: 1.6 (Sol Conseil 2022, Nyon, Switzerland)

Preliminary crop was clover, followed by potatoes.

For the design, varieties were arranged in a 6 × 6 lattice design with four replications. The experimental unit was a plot of 9 m^2^ (1.5 m in row and 6 m in range direction including paths and wheal tracks), resulting in a harvested surface of 7.05 m^2^.

Wheat was drill-sown in 8 rows per plot with a row distance of 0.125 m on November 12, 2019. Sowing density was 350 plants m^-2^.

Additional management information:

20.01.2020 165 kg/ha Landor 0/20/30

17.03.2020 368 kg/ha Nitrate magnésien soufré 24/0/0 (88 kg N/ha)

18.03.2020 0.5 kg/ha Artist + 1 l/ha Netzmittel + 200 g/ha Othello Star

04.05.2020 294 kg Sulfonitrate 26/0/0 (77 kg N/ha)

27.02.2020 Harvest

(Total 165 kg N/ha)

### Site FIP

The site FIP is located at the ETH research station of agricultural sciences in Lindau Eschikon, Switzerland (47.449 N, 8.682 E, 556 m a.s.l.). The soil type is an eutric cambisol consisting of 21% clay and 21% silt. Organic matter content is 3.5% and pH 6.7. Soil characteristic were determined in 2015 (Eric Schweizer AG, Thun, Switzerland). Preliminary to wheat (*Triticum aestivum L*.), soybeans (*Glycine max* (L.) Merr.) and buckwheat (*Fagopyrum esculentum* Moench) were grown. After preliminary crops were harvested, the soil was plowed and harrowed before wheat was drillsown.

#### FIP19

For the design, 36 test varieties were arranged in four replications equally allocated across two lots of the FIP. The experimental unit was a plot of 6.1 m^2^ (1.5 m in row and 6 m in range direction including paths and wheal tracks). The varieties were allocated in a row-column design as follows: Full replicates in row direction consisted of six rows by six ranges. Blocks in range direction consisted in two ranges spanning the 24 rows of both lots, thus holding 1.33 replications per genotype. This dimension was introduced to cover the spatial trend of the sloped field in an upper, central and lower part. The design was generated using the R-package DiGGER.

Wheat was sown in 9 rows per plot with a row length of 5 m and a row distance of 0.125 m on October 17, 2018. Sowing density was 400 plants m^-2^. One day after sowing, herbicide (Herold SC, Bayer AG, Leverkusen, Germany) was applied to ensure weed free plots. Several fungicides and insecticides were applied in spring to ensure healthy plants.

On February 27, 2019, April 8 and May 27, 52 kg N ^-1^, 72 kg N ^-1^ and 24 kg N ha^-1^ were applied. Additionally, in total 92 kg phosphorous ha^-1^, 120 kg potassium ha^-1^ and 15 kg magnesium ha^-1^ were applied.

#### FIP20

For the design, 36 test varieties were arranged in three replications on one lot of the FIP. The experimental unit was a plot of 6.1 m^2^ (1.5 m in row and 6 m in range direction including paths and wheal tracks). The varieties were allocated in a row-column design as follows: Full replicates in row direction consisted of six rows by six ranges. Blocks in range direction consisted in two ranges spanning the 18 rows of the lot. This dimension was introduced to cover the spatial trend of the sloped field in an upper, central and lower part. The design was generated using the R-package DiGGER.

Wheat was sown in 9 rows per plot with a row length of 5 m and a row distance of 0.125 m on October 17, 2019. Sowing density was 400 plants m^-2^. Eight days after sowing, herbicide (Herold SC, Bayer AG, Leverkusen, Germany) was applied to ensure weed free plots. Several fungicides and insecticides were applied in spring to ensure healthy plants.

On March 3, 2020, 26 kg N ha^-1^ and 58 kg Mg ha^-1^ were applied. On April 27, 2020, 72 kg ha^-1^ and 15 kg Mg ha^-1^ were applied. On May 7, 2020, 24 kg N ^-1^ and 5 kg Mg ha^-1^ were applied.

### Site Strickhof

The site Strickhof is located at the Strickhof Landwirtschaftsschule in Lindau Eschikon, Switzerland (47.445 N, 8.683 E, 535 m a.s.l.). The soil was a skeleton rich Cambisol (Landwirtschaftsamt des Kanon Zürich, 1992) with 34% clay, 3.3% organic matter and a pH of 7.4 (Martin Bertschi, Strickhof, personal correspondence, October 2019).

#### Strickhof19

There were four replications on one lot at the Strickhof site where the genotypes were arranged in a random complete block design. The management was less intensive, following the guidelines of Federal Swiss pesticide reduction program “Extenso”. No insecticides, fungicides or plant growth regulators were applied. Fertilization regime was following common agricultural practice. Sowing took place on October 17, 2018, with the same specifications as at ETH-FIP site but at a length of about 6.4 m.

## Footnotes

1 Please note our consciously chosen relaxed definition of phenomic selection (PS) in contrast to Rincent et al. (2018) and Robert et al. (2022): While Rincent et al. (2018) restricted PS to depend on similar statistical models as GS, Robert et al. (2022) went one step further and defined the input of PS to be based on near-infrared (NIR) spectroscopy data. Unlike NIRs, however, most phenomics’ techniques allow resolving environmental-specific phenotypes, calling for other statistical models than GS, and widening the scope of PS far beyound NIRs data.

